# Heterogeneity of Tethered Agonist Signaling in Adhesion G Protein-Coupled Receptors

**DOI:** 10.1101/2022.07.14.500097

**Authors:** Andrew N. Dates, Daniel T.D. Jones, Jeffrey S. Smith, Meredith A. Skiba, Maria F. Rich, Maggie M. Burruss, Andrew C. Kruse, Stephen C. Blacklow

**Affiliations:** Department of Biological Chemistry and Molecular Pharmacology, Blavatnik Institute, Harvard Medical School, Boston, MA 02115, USA; Department of Dermatology, Brigham and Women’s Hospital, 221 Longwood Avenue, Boston, MA 02115; University of Cincinnati School of Medicine, Department of Molecular Genetics, Biochemistry, and Microbiology, Cincinnati, OH 45267, USA; Department of Cancer Biology, Dana Farber Cancer Institute, Boston, MA 02215, USA

## Abstract

Adhesion G Protein-Coupled Receptor (aGPCR) signaling influences development and homeostasis in a wide range of tissues. In the current model for aGPCR signaling, ligand binding liberates or unmasks a highly conserved tethered agonist (TA) that acts as an intramolecular ligand to stimulate G protein coupling. However, a systematic, comprehensive examination of signaling has not been performed for all aGPCR family members. Here, we report a platform for profiling TA-dependent activities of aGPCRs in several different assays and apply it to all 33 human family members. Activity profiling identified a heterogeneity of responses among these aGPCRs, with only ∼50% showing robust, TA-dependent signals. AlphaFold2 predictions assessing TA engagement in the predicted intramolecular binding pocket aligned with the TA-dependence of the cellular responses. The signaling information in this dataset is a comprehensive resource relevant for the investigation of all human aGPCRs and for targeting aGPCRs therapeutically.

## Introduction

As the largest class of membrane proteins in the human genome, G protein-coupled receptors (GPCRs) are central modulators of physiology^1^. The 33 Adhesion G Protein-Coupled Receptors (aGPCRs, GPCR family B2) comprise the second-largest family of GPCRs, and they influence development, cell maturation, and tissue maintenance through roles in synapse formation, myelination, immune cell migration, auditory perception, and other critical processes^2–9^

Despite their abundance in the human genome, aGPCRs are relatively understudied compared to other GPCRs^10–14^. Unlike GPCRs from other families, which have been systematically screened for coupling to G proteins and β-arrestins^15–19^, aGPCRs have not been comprehensively profiled for their signaling properties, limiting efforts to link molecular mechanism to physiological role.

The aGPCRs are distinguished from other GPCRs by their modular domain organization, which has three distinct elements (Fig. 1A). At the N-terminus, there is an adhesion module that typically consists of a series of protein interaction domains responsible for ligand binding^20^. Between the adhesion module and the membrane lies a regulatory domain, called the GPCR autoproteolysis-inducing (GAIN) domain. The third module consists of the seven-transmembrane (7TM) GPCR domain, which couples to intracellular effectors to transduce signals across the membrane. The aGPCRs also have cytoplasmic C-terminal tails of variable length.

**Fig. 1.**
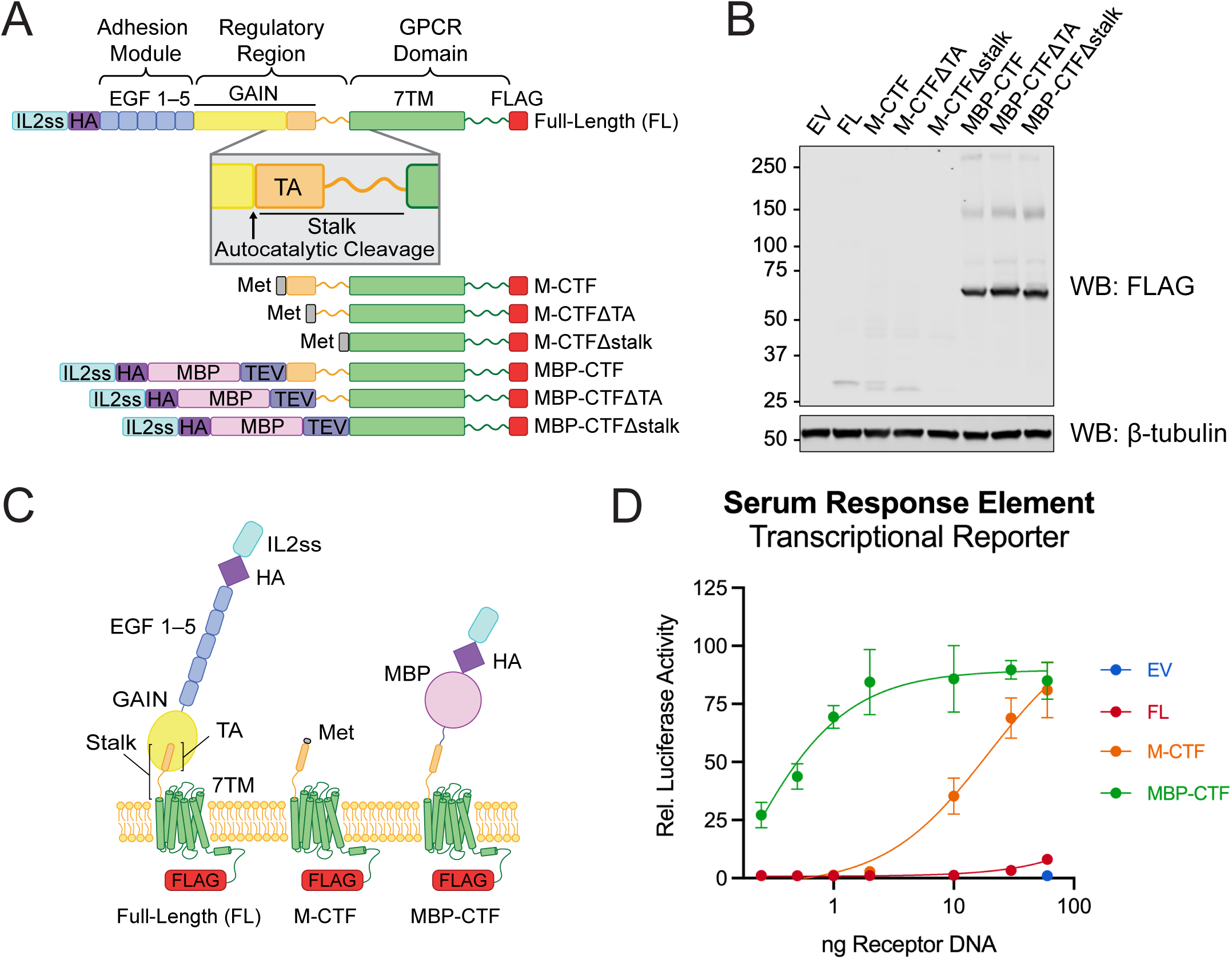
– A fusion protein system to evaluate aGPCR signaling. (A) Domain architectures of constructs used to evaluate signaling in ADGRE5. In the full-length (FL) protein, autocatalytic cleavage occurs within the GAIN domain, immediately prior to the TA peptide, which is sequestered within the folded GAIN domain. C-terminal fragment proteins with exposed TA peptides are expressed with an N-terminal methionine (M-CTF) or after a signal sequence, HA epitope tag, and maltose-binding protein fusion domain (MBP-CTF). The design of the ΔTA and Δstalk variants is also shown. (B) Western blot detection of the FLAG C-terminal epitope tag to assess total protein abundance, with β-tubulin as a protein loading control. (C) Schematic diagrams of ADGRE5 signaling constructs. (D) Comparison of ADGRE5 variants in the SRE transcriptional reporter signaling assay as a function of the amount of transfected DNA. All data points are normalized to the mean value for empty vector (60 ng). Dose-response curves were fit for each titration curve using nonlinear regression. Data are presented as mean ± SEM of three independent biological replicates (n=3).

A distinctive feature of some GAIN domains is that they can undergo autocatalytic cleavage during maturation^21,22^. This self-processing step divides these aGPCRs into two non-covalently associated subunits: an N-terminal fragment (NTF) containing the adhesion module and most of the GAIN domain and a C-terminal fragment (CTF) comprised of the residual “stalk” sequence from the GAIN domain, the 7TM domain, and the cytoplasmic tail (Fig. 1A). The stalk is roughly 15-25 residues long, containing a conserved N-terminal segment of seven amino acids that is largely hydrophobic. The first seven stalk residues are sequestered as a β-strand in the GAIN domain under resting conditions^21,23–25^. For several aGPCRs, this sequence can be liberated from the GAIN domain, putatively in response to force delivered to the NTF by bound ligand, to function as a tethered agonist (TA) for signal activation^14,26–33^.

An atomic-level view of tethered agonism in aGPCRs has emerged from single particle cryo-electron microscopy (cryo-EM) studies of seven CTFs from different aGPCRs in G protein-bound states^34–38^. For these aGPCRs, the structural studies define a conserved binding pose of the TA in the orthosteric site, which facilitate transition of the GPCR to an active, G protein-bound state.

Although the TA-induced signaling model clearly applies for some family members, including those for which active state structures have been reported, whether TA ligation elicits a signaling response for all aGPCRs remains unclear. For about half of aGPCRs, TA-dependent signaling has not been established, in part because of the difficulty in developing an assay generally applicable to all aGPCRs. Of the aGPCRs shown to exhibit TA-dependent signaling, few have been evaluated side-by-side and even fewer across a comprehensive set of functional readouts such as signal-induced transcription, G protein recruitment, and receptor internalization, making it difficult to compare signaling properties among them^11–13^.

Here, we report a platform for profiling TA-dependent activities of aGPCRs in several different assays and apply it to all 33 human family members. Activity profiling identified a heterogeneity of responses, with only ∼50% of family members showing robust, TA-dependent signals, largely consistent with AlphaFold2 predictions of TA engagement^39,40^. The signaling information in this dataset should be broadly relevant for tool compound discovery, therapeutic development, and future *in vivo* studies.

## Results

### Development of an assay platform for assessment of aGPCR signaling

We selected ADGRE5 (also known as CD97) as a prototype for development of a general approach to evaluate aGPCR signaling^41–43^. We expressed recombinant full-length protein (FL), a C-terminal fragment (CTF) with an N-terminal methionine residue (methionine-capped, M-CTF), and two truncated forms of the CTF, one lacking the seven-residue tethered agonist sequence (M-CTFΔTA), and another lacking all fifteen residues preceding TM1 (a “stalkless” construct called M-CTFΔstalk), in HEK293 cells (Fig. 1A). The steady state amounts of these methionine-capped proteins varied substantially (Fig. 1B), however, making it difficult to compare signaling activities of the variants produced using this approach.

We used protein engineering to achieve consistency in expression among ADGRE5 variants. Active state structures of CTFs of aGPCRs show that the TA is positioned in a conformation with an exit vector compatible with an N-terminal fusion extending into the extracellular space (Fig. S1A)^34–38^. We postulated that insertion of a stable N-terminal fusion protein would permit TA-dependent signaling and reduce variation in steady-state protein abundance among different aGPCRs and their variants, enabling the measurement of TA-dependent and TA-independent signaling activity for each aGPCR using a single expression cassette. We designed the same series of ADGRE5 variants with a secretion signal sequence at the N-terminus, followed by an HA epitope tag, the thermostable maltose binding protein (MBP), and a TEV protease cleavage site preceding CTF (MBP-CTF), CTFΔTA (MBP-CTFΔTA), and CTFΔstalk (MBP-CTFΔstalk) sequences (Fig. 1A,C). We compared the steady state amounts of these proteins in HEK293 cells to one another and to the methionine-capped CTFs (Fig. 1B). Unlike the M-CTF proteins, the MBP fusion proteins were present at similar – and substantially greater – total protein abundance when transfected in equivalent amounts, enabling their direct comparison in downstream functional assays.

We compared the signaling properties of these ADGRE5 variants in a transcriptional reporter assay, an established and sensitive metric of GPCR activity^44^. Transfection of full-length ADGRE5 into HEK293 cells resulted in low basal activity of the Serum Response Element (SRE) reporter, whereas transfection of either MBP-CTF or M-CTF induced a strong, saturable SRE reporter signal (Fig. 1D). Strikingly, the MBP-CTF fusion was roughly 30-fold more potent than M-CTF as a function of plasmid dose without the need for addition of TEV to catalyze release of the N-terminal fusion domain. Although the MBP-CTF fusion was produced in greater amounts than the M-CTF protein when equivalent quantities of DNA were transfected (Fig. S1B), this difference in abundance does not fully account for its increased potency. In contrast, the MBP-CTFΔTA and MBP-CTFΔstalk proteins show negligible signaling activity, confirming that the stalk sequence is necessary to induce a transcriptional reporter signal (Fig. S1C). The identity of the fusion domain also had little effect on the signaling activity of ADGRE5, because a range of different N-terminal fusion sequences also led to constitutive signaling by the CTF (Fig. S1D,E).

We serially deleted linker residues to determine the minimal spacer length between the fusion protein and the TA that permitted signaling. Deletion of the entire TEV site and even the first serine residue (S531^P1’^) of the tethered agonist from the ADGRE5 MBP-CTF protein did not silence its constitutive signaling activity. Additional removal of S532^P2’^, however, reduced the signaling activity to that of the quiescent FL receptor (Fig. S2). This decrease in activity resulted from steric interference by the MBP fusion, because reinsertion of the TEV cleavage site between MBP and F533^P3’^ restored activity to that of the parental MBP-TEV-CTF protein (Fig. S2).

We analyzed the effects of systematically mutating individual ADGRE5 TA residues on reporter gene activity to interrogate structure-function relationships in the ADGRE5 stalk. Our scanning mutagenesis showed that signaling from ADGRE5 was intolerant of either alanine or lysine substitution at both F533 (P3’) and L536 (P6’) (Fig. S3A), consistent with the known importance of these positions for signaling by other aGPCRs and the deep burial of these TA positions in active state aGPCR structures^26,27,34–38^. The observed effects of other stalk sequence mutations aligned with a homology model of the ADGRE5 CTF (Fig. 3B)^40^, with solvent exposed and predicted linker residues tolerant of both alanine and lysine substitutions, and other partially buried positions tolerant only of alanine (or glycine, for P4’ residue A534 and P8’ residue A538). The MBP-CTF platform resulted in similar amounts of expressed protein for all mutants tested except F533A and L536A, both of which were signaling incompetent in the reporter assay and present in lower abundance than wild-type protein at steady state as judged by western blot (Fig. S3C,D). The reduced abundance of these variants is likely a consequence, rather than a cause, of their signaling incompetence; the analogous signaling-deficient F569A mutant in the TA of ADGRF1 was also nearly undetectable when expressed as a CTF protein lacking a stabilizing domain such as MBP^45^.

### Assessment of TA-dependent signaling across the complete human aGPCR family

We evaluated the ability of each human aGPCR to be activated by its TA peptide in the context of an HA-MBP-TEV-CTF-FLAG fusion analogous to the ADGRE5 prototype. We aligned the GAIN domain sequences of all human aGPCRs to identify the P1’ position of each and assembled a complete set of CTF fusion proteins with the P1’ residue positioned immediately after the TEV site (designated MBP-CTF) (Fig. S4). We also made a matched set of fusion proteins lacking the tethered agonist in which the conserved P1’-P7’ residues that comprise the TA were deleted, designating these variants MBP-CTFΔTA (Fig. S4). For ADGRA1, which lacks a GAIN domain, the amino acid sequence of the full-length receptor was present in the MBP-CTF construct, and we deleted all residues prior to the start of TM1 in the MBP-CTFΔTA construct. All MBP-CTF and MBP-CTFΔΤΑ proteins were readily detectable by western blot when expressed in HEK293T cells except for ADGRE4, which is a predicted pseudogene (Fig. S5). Comparable amounts of protein were expressed by almost all matched MBP-CTF and MBP-CTFΔΤΑ protein pairs, showing that the MBP fusion strategy suppressed variability in expression not merely for ADGRE5, but for the full family of aGPCRs.

We assessed TA-dependent and TA-independent signaling activity by measuring the ability of each MBP-CTF and MBP-CTFΔΤΑ protein to induce the expression of a luciferase reporter gene under the control of four different response elements^44^. Assays were performed using a serum response element (SRE) reporter, a cAMP Response Element (CRE) reporter, a Nuclear Factor of Activated T Cells Response Element (NFAT-RE) reporter, and a Serum Response Factor Response Element (SRF-RE) reporter (Fig. 2A). Signaling patterns varied substantially among the different subfamilies, and there was even heterogeneity of response patterns within the different subfamilies (Fig. 2B). We note that the SRE and SRF-RE reporters showed near identical response patterns across aGPCR family members, suggesting that they are integrating very similar signaling inputs. Of the 32 family members, 16 showed robust TA-dependent transcriptional responses with at least one reporter, including proteins from each of the D, E, F, G, and L subfamilies. However, no proteins of the A, B, C, or V subfamilies showed TA-dependent expression of the luciferase reporter, nor did ADGRE2, ADGRE3, ADGRF2, ADGRF5, ADGRL1, or ADGRL4 show a TA-dependent reporter response. Members of the B, C, and V subfamilies, as well as ADGRG3 did produce activity with some reporters, but this transcriptional response was independent of the presence of the TA. The TA-independent activity was also typically substantially weaker than the TA-dependent signal observed in D, E, F, G, and L family members. Together, these transcriptional reporter data define a functional signaling landscape for the complete family of human aGPCRs, identifying a spectrum of TA responsiveness among them.

**Fig. 2.**
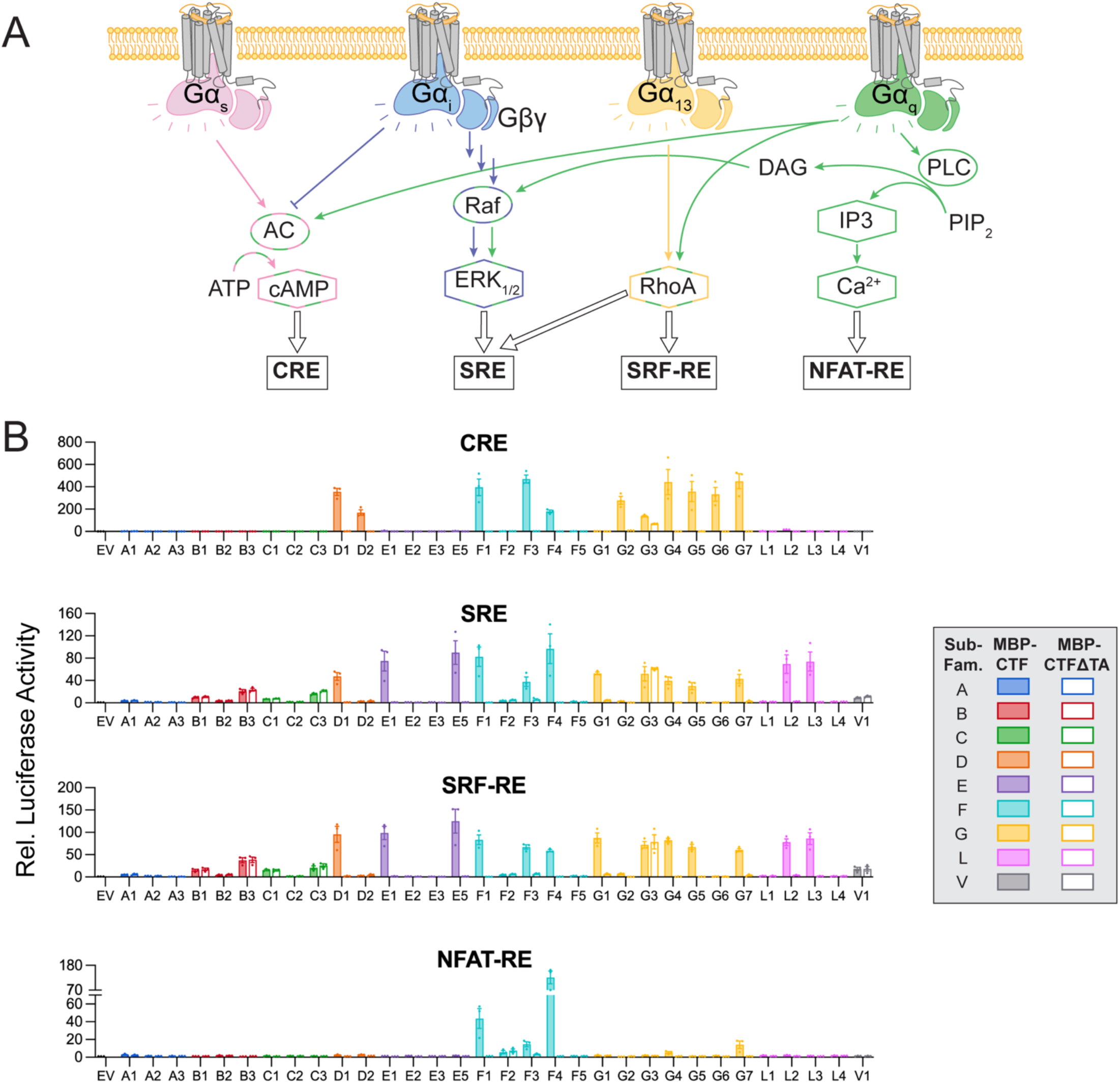
– Transcriptional reporter responses for all human aGPCR family members. (A) CRE, SRE, SRF-RE, and NFAT-RE transcriptional reporter gene assays, illustrating potential sources of stimulation downstream of G protein signaling. (B) CRE, SRE, SRF-RE, and NFAT-RE reporter activity of cells expressing either MBP-CTF (filled bars) or MBP-CTFΔTA (open bars). Data are normalized to the mean value for empty vector and are presented as mean ± SEM of three independent biological replicates (n=3).

Transcriptional activity is a distal signaling readout downstream of G proteins, which are the primary transducers of GPCR-initiated signaling cascades. We also adapted a series of TRUPATH G protein biosensors to measure the TA-dependent ability of each aGPCR to stimulate the dissociation of Gα subunits from Gβγ complexes, a direct consequence of coupling between the aGPCR and the G protein^46^. These assays detect changes in Bioluminescence Resonance Energy Transfer (BRET) associated with the dissociation of RLucII-fused Gα subunits from GFP-fused Gβγ complexes upon receptor activation (Fig. 3A). For GPCRs activated by *trans* addition of a soluble agonist, biosensor activation is typically reported as a ligand-dependent change in BRET efficiency (ΔBRET). Because the TA is an intramolecular aGPCR ligand, here we measured the BRET efficiencies of the MBP-CTF and MBP-CTFΔTA (Fig. S6) proteins and calculated the difference between them to determine the TA-dependent change in BRET efficiency (ΔBRET). The aGPCRs showed distinct patterns of TA-dependent G protein activation with the G_s_, G_q_, G_i1_, and G_13_ biosensors (Fig. 3B–E). The TA-dependence of aGPCR signaling activity in the BRET assay closely tracked the aGPCR activity in the transcriptional reporter assay, with proteins of the A, B, C, and V subfamilies again quiescent in the BRET assay. Strikingly, the G_13_ BRET biosensor was the most frequently activated, indicating a G protein preference of aGPCRs that differs from family A and family B1 GPCRs (Fig. 3F). Together, the combination of reporter and BRET assays revealed that the aGPCRs fall into two distinct groups: those that transduce TA-dependent signals and those that do not.

**Fig. 3.**
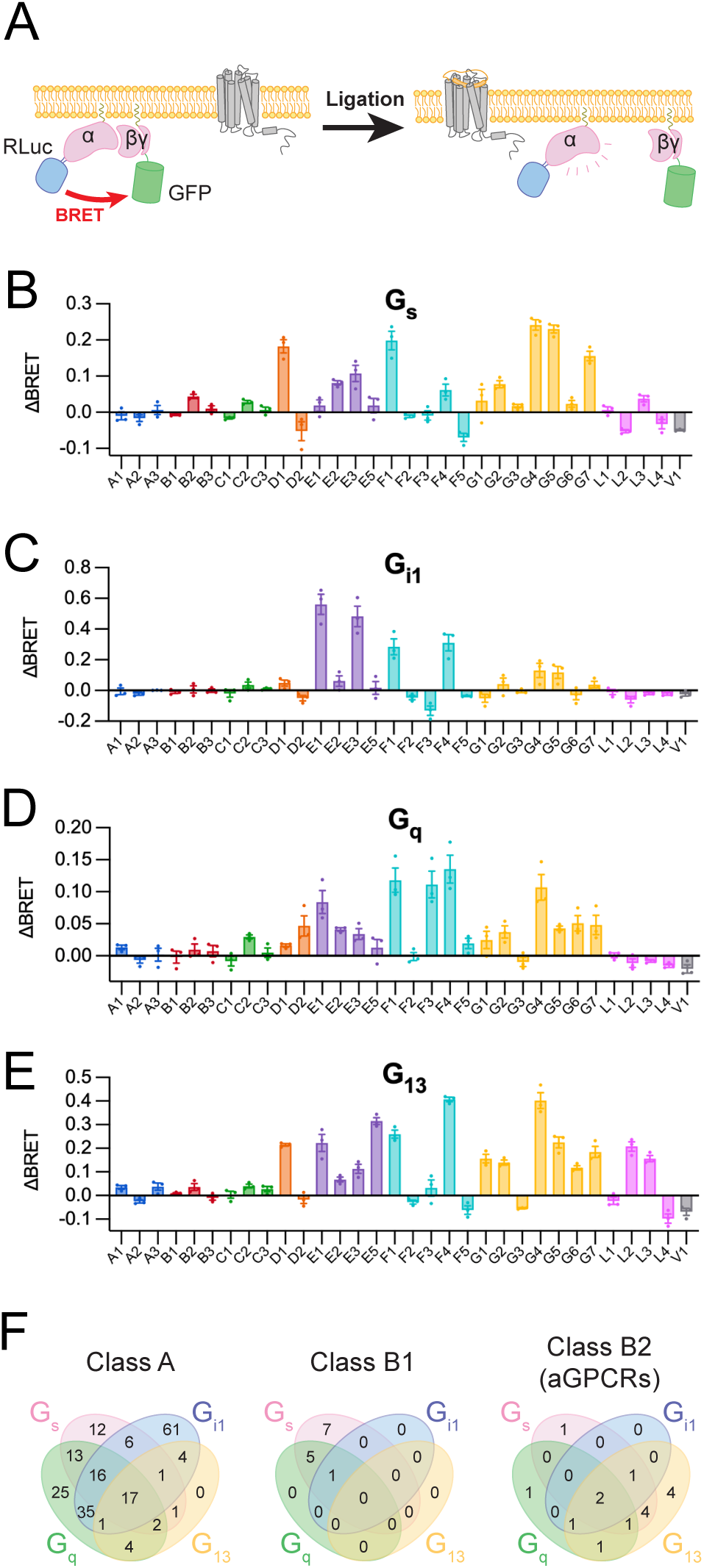
– Tethered agonist-dependent G protein activation activity of all human aGPCRs. (A) Schematic depicting G protein biosensor assays, which measure the dissociation of Gα subunits from Gβγ complexes. (B–E) TA-dependent activation of the G_s_ (B), G_i1_ (C), G_q_ (D), and G_13_ (E) biosensors for all aGPCRs. ΔBRET values are the difference in resting BRET ratios between MBP-CTF and MBP-CTFΔTA for each protein pair. Data are presented as mean ± SEM of three independent biological replicates (n=3). (F) Summaries of G protein biosensor activation for family A^19^, B1^19^, and B2 GPCRs. Family B2 signals were assigned using one-way ANOVA with Tukey’s multiple-comparison post-hoc test to compare ΔBRET values for each receptor with that of EV (zero) (* P<0.05).

### Assessment of aGPCR localization

We also measured the steady state cell-surface abundance of all MBP-CTF and MBP-CTFΔTA matched pairs by flow cytometry, using an anti-HA antibody recognizing the extracellular part of the protein. Whereas western blotting showed that the amount of expressed protein was roughly equivalent for each MBP-CTF and MBP-CTFΔTA matched pair (Fig. S5), many of the MBP-CTF proteins from the D, E, F, G and L families were depleted from the cell surface when compared with their MBP-CTFΔTA counterparts (Fig. 4). The cohort of proteins that showed reduced MBP-CTF staining relative to their matched MBP-CTFΔTA partners closely correlated with the group of proteins that showed TA-dependent activity in the reporter gene and G protein biosensor assays. The tendency of autonomously active, MBP-CTF forms of TA-dependent aGPCRs to be depleted from the cell surface relative to their inactive, MBP-CTFΔTA forms is consistent with examples from other families of GPCRs in which receptor activation is followed by receptor internalization^47^.

**Fig. 4.**
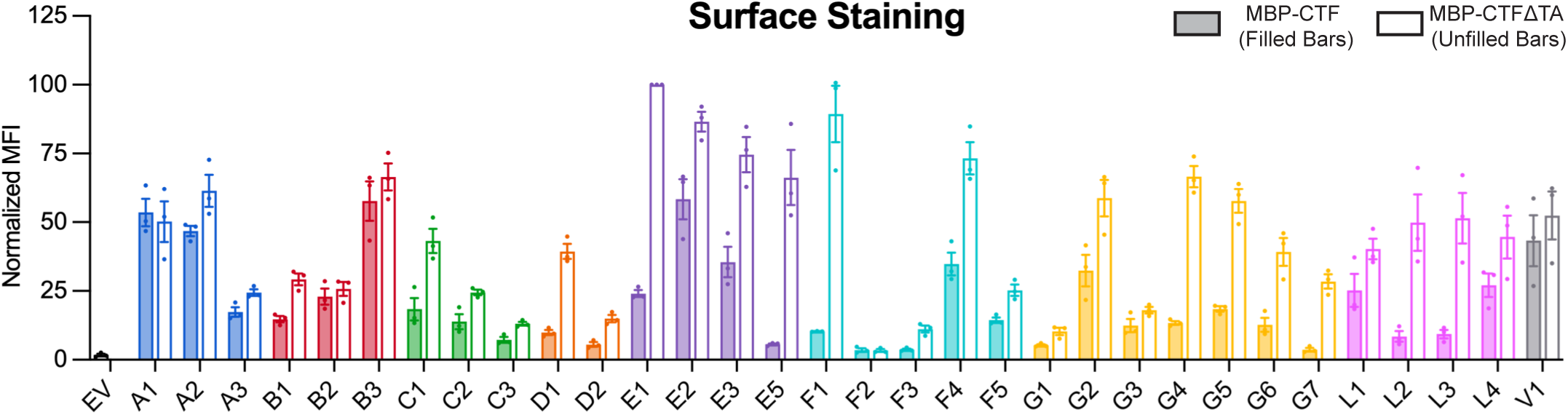
– Cell surface amounts of all aGPCR protein constructs. α-HA surface staining of non-permeabilized HEK293T cells expressing HA-MBP-CTF-FLAG (filled bars) and HA-MBP-CTFΔTA-FLAG (open bars) proteins. Data from three biological replicates (n=3) are presented as mean fluorescence intensity (MFI) ± SEM, normalized to the value for ADGRE1, which was set to 100.

### Comparison of structural predictions with signaling and internalization data

We modeled the structure of each CTF using Colabfold^40^ to assess whether the TA sequence was predicted to be bound intramolecularly for each CTF (*i.e.*, without the MBP fusion domain). The predicted structures consistently adopted one of two conformations (Fig. 5A, Fig. S7), either with the TA occupying the orthosteric site of the 7TM domain and a kink in the TM6 helix at residue G^6.50^, or with the TA unbound and an unkinked TM6 helix. Strikingly, the structural predictions closely correlated with the TA-mediated signaling responses from our transcriptional readouts, G protein coupling assays, and surface staining analyses, providing a functional landscape of aGPCR activities extending from agonist binding to receptor desensitization (Fig. 5B-F).

**Fig. 5.**
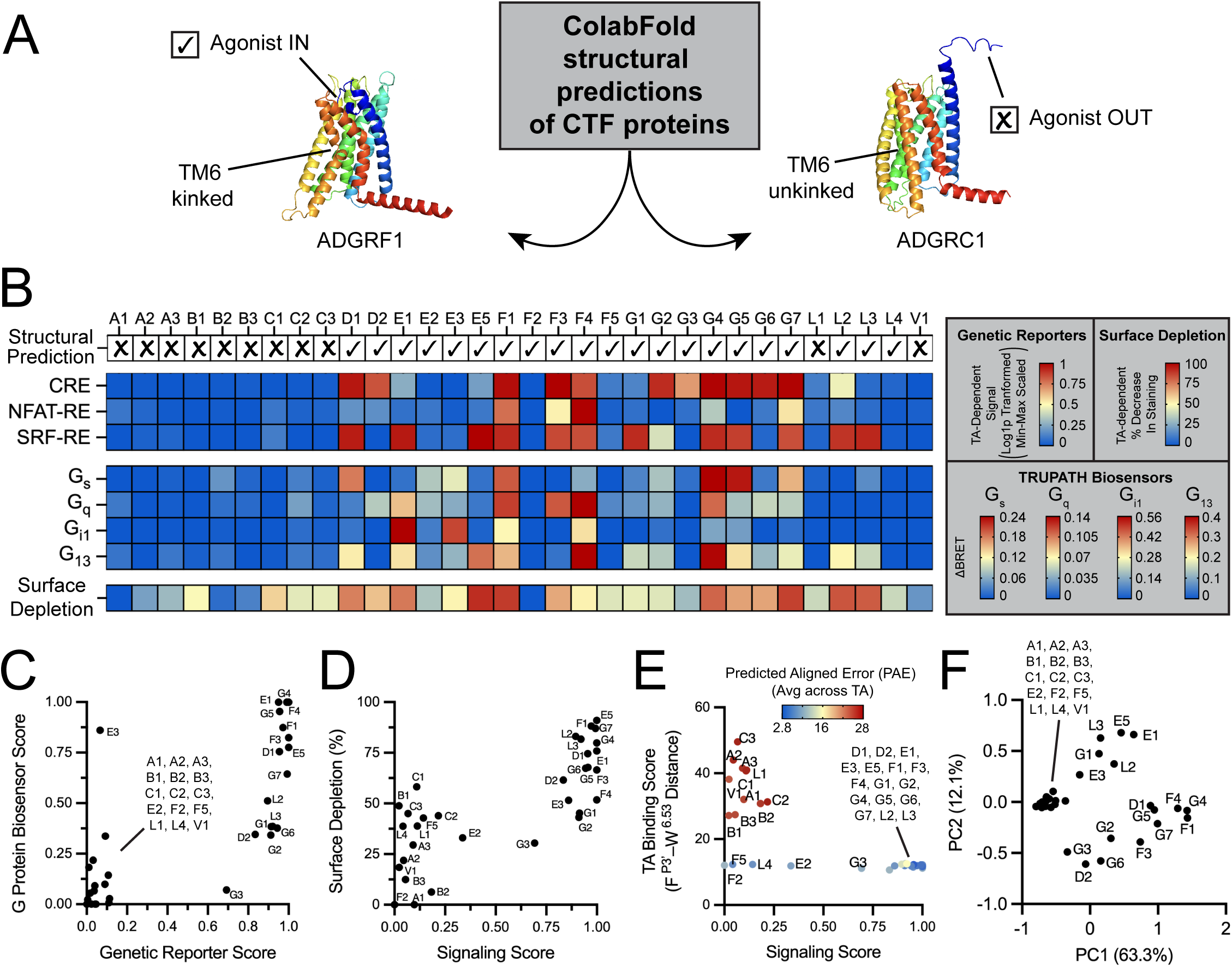
– Structural predictions and heat map representations of responses for all human aGPCRs. (A) Representative examples of CTF “in” and “out” conformations predicted by ColabFold, showing the position of the TA peptide and degree of kinking by TM6, which is a hallmark of Family B GPCR activation. (B) Heat map summarizing TA-dependent transcriptional activation, G protein activation, surface localization, and ColabFold predictions for the human aGPCRs. (C–E) Scatter plots correlating G protein biosensor score with genetic reporter score (C), surface depletion with signaling score (including all reporters and G protein biosensors) (D), and ColabFold-generated TA binding scores with signaling score (E) for each aGPCR. Points in (E) are colored by the Predicted Aligned Error confidence metric averaged across the seven TA residues for each receptor. (F) Principal Component Analysis of all measured TA-dependent responses when each measured response is scaled zero to one across all aGPCRs evaluated in this study.

Despite the strong conservation of the TA sequence and autoproteolysis motif (Fig. S8), the signaling properties of the human aGPCRs are heterogeneous with respect to their TA dependence. The invariance of the TA sequence thus suggests that structural motifs within the 7TM itself accounts for the activity differences. Multiple sequence alignment revealed that residue G^6.50^ within the 7TM domain is completely conserved in TA-activated aGPCRs but largely absent in the quiescent receptors (Fig. 6A). This residue lies in a PXXG motif also found in family B1 (Secretin) GPCRs^48^, which is required to facilitate the kinking of the TM6 helix characteristic of the fully active state in family B GPCRs. Mutation of this position to alanine in four TA-activated aGPCRs (ADGE5, ADGRF1, ADGRG1, and ADGRL3) impaired their ability to induce genetic reporters, confirming the importance of this residue in TA-dependent aGPCR activation (Fig. 6B, Fig. S9), and offering a molecular rationale for the observed functional heterogeneity of the human aGPCRs.

**Fig. 6.**
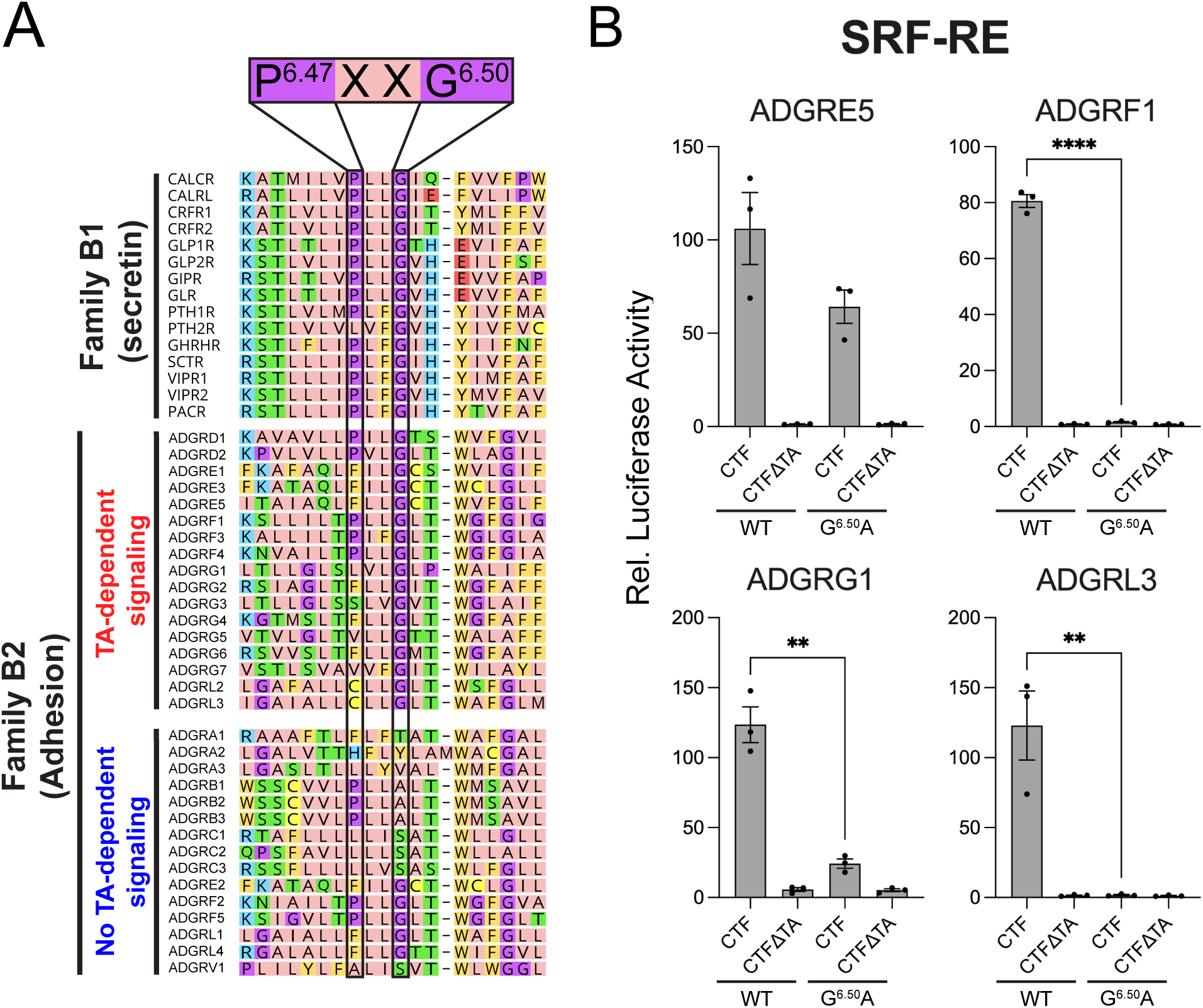
– Conservation and functional relevance of the PXXG activation motif in aGPCRs. (A) Multiple sequence alignment of TM6 across human Family B GPCRs, with aGPCRs segregated by TA-signaling competence in our MBP fusion system. (B) Effects of alanine substitution at G^6.50^ within the PXXG motif of four TA-activated aGPCRs in the SRF-RE reporter. Data are normalized to the mean value for empty vector and are presented as mean ± SEM of three independent biological replicates (n=3). Unpaired two-tailed T-tests were performed to compare WT and mutant CTF proteins (** P<0.01, **** P<0.0001).

It is possible that, for certain receptors, the N-terminal MBP fusion interfered with access of the TA to its putative binding site on the 7TM domain, preventing TA-dependent signaling. We evaluated this possibility by testing whether TEV protease-catalyzed release of the MBP fusion domain would restore TA-dependent signaling activity for inactive aGPCRs. Co-expression of a secreted, ER-retained form of TEV (secTEV)^49^ with three TA-activated receptors (ADGRE5, ADGRF1, and ADGRL3) led to processing of each MBP-CTF protein at the TEV cleavage site (Fig. S10) and did not substantially change the signaling activity of these proteins in the SRF-RE reporter assay. Co-expression of secTEV also led to MBP release for all receptors that were silent in the TA-dependent signaling assays, except ADGRG3, and yet none of these receptors exhibited TA-dependent signaling in the CRE, SRF-RE, and NFAT-RE reporters. These data confirm that the quiescent subset of aGPCRs do not exhibit TA-dependent signals, even when the TA is completely exposed at the N-terminus after MBP removal (Fig. S10).

## Discussion

Numerous GPCRs belonging to families A, B1, and C have been profiled to assess G protein coupling activity and downstream transcriptional responses, but platform-based profiling of aGPCRs has been elusive because the agonist is an intramolecular ligand that is masked in the mature full-length protein. By constructing an MBP fusion platform that was modular and generalizable, we were able to evaluate the coupling and signaling response of each aGPCR to its intramolecular TA.

We compared the signaling activities of two variants for each receptor, one with and one without the TA (referred to as CTF and CTFΔTA, respectively), in reporter gene assays and with G protein biosensors, and used ColabFold to predict whether or not the TA would be bound in the orthosteric site. Our findings uncovered a striking and unexpected dichotomy among the aGPCRs, in which some exhibited TA-dependent signals and some did not.

Our studies revealed tethered agonism for three aGPCRs that had not previously been evaluated: ADGRD2, ADGRE1, and ADGRE3. We also confirmed G protein coupling responses for fourteen other receptors and identified secondary signaling responses for ADGRD1, ADGRG2, ADGRG6, ADGRF4, and ADGRL2. The aGPCRs of the A, B, C, and V subfamilies, however, as well as ADGRE2, ADGRF2, ADGRF5, ADGRL1, and ADGRL4, did not show evidence of TA-dependent signaling in transcriptional reporter assays or with G protein biosensors. ColabFold predictions of CTF structures were also consistent with the experimental signaling data with few exceptions.

It is possible that some of the receptors failing to exhibit TA-dependent signaling in our system may signal through the TA *in vivo*. ADGRE2, for example, can signal through G_16_ heterotrimers^50^, which are not present in HEK293T cells, and the ADGRF2 proteins used in this study were poorly expressed. AlphaFold2 predicted these two proteins, as well as ADGRF5 and ADGRL4, to be TA binding-competent, and these receptors contain an intact PXXG motif, but these proteins showed no TA-dependent signaling in our assays. Nevertheless, release of the MBP to reveal a free TA sequence at the N-terminal end of the CTF for all of the receptors that did not exhibit TA-dependent signaling did not produce any TA-dependent signals in the reporter gene assays.

Some of the receptors that did not produce TA-dependent signals did have increased basal amounts of TA-independent activity in some of the reporter gene assays. Whether this activity represents functionally important basal signaling that is biologically relevant independent of ligand induction remains unclear.

We used flow cytometry to assess the cell-surface amounts of each protein because receptor internalization can be a mechanism for desensitizing or compartmentalizing GPCR signaling^47,51,52^. The CTFΔTA form of the protein tended to be present in higher amounts at the cell surface than the CTF form for proteins that exhibited TA-dependent signals, suggesting an agonist-dependent effect on aGPCR internalization. These data also suggest that the constitutive signaling properties of CTF proteins may complicate the use of surface abundance to normalize for the relative potency of different proteins in activity assays.

Heterogeneity of TA-dependent signaling among the aGPCRs existed despite the high conservation of the TA sequence. Its high conservation, however, may derive from its role as the terminal β-strand within the core of the GAIN domain in each receptor^21^. The segregation of the aGPCRs into two groups may instead be due to the presence or absence of the TM6 helix PXXG motif, which is critical for the activation of family B1 GPCRs^48^.

Although autocatalytic cleavage in the GAIN domain is necessary for dissociative presentation of the TA sequence at the N-terminus of the CTF, self-cleavage within the GAIN domain does not occur in all aGPCRs. Indeed, receptors lacking canonical cleavage motifs, including ADGRE1 and ADGRF4, exhibited TA-dependent signaling. In ADGRF4, a polybasic consensus sequence for furin cleavage lies ∼30 residues toward the N-terminus from the TA and may substitute for the autocatalytic cleavage event to enable subunit dissociation. For ADGRE1, however, no such alternative cleavage site exists, consistent with the possibility that cleavage-independent access of the TA sequence can occur, as has been proposed^53,54^.

Finally, the aGPCRs showed a general coupling preference for G_12/13_ heterotrimers, the least frequently activated effectors for other GPCR families. G_12/13_ activation leads to actin cytoskeletal rearrangement^55,56^, which correlates with the biological processes governed by many aGPCRs. Our catalog of TA-dependent signals and their effector dependencies should be a valuable resource to assist in analyzing the molecular bases and effector dependencies of aGPCR-driven biological phenotypes, and ultimately should facilitate the development of tool compounds for therapeutic development.

## Acknowledgments

This work was supported by a grant from the Warren Alpert foundation (to S.C.B.) and award 1 R35-CA220340 (to S.C.B). A.N.D. was supported by a fellowship from the Fujifilm Corporation. M.A.S. was supported by a Merck Postdoctoral Fellowship from the Helen Hay Whitney Foundation. J.S.S. is supported by National Institutes of Health Dermatology Training Grant 5T32AR007098-49 (J.S.S.) and the Dermatology Foundation (J.S.S.). We thank Justin English (University of Utah) for generously providing the tricistronic TRUPATH plasmids. We also thank members of the Blacklow and Kruse labs for helpful discussions.

## Author Contributions

A.N.D. and S.C.B. conceived of the research. A.N.D., D.T.D.J., J.S.S, M.A.S, A.C.K, and S.C.B designed experiments. A.N.D., M.F.R., and M.M.B. performed experiments. A.N.D., D.T.D.J., J.S.S., M.A.S, A.C.K., and S.C.B. interpreted data. A.N.D. and S.C.B. wrote the manuscript with input from all authors.

## Methods

### Materials

DMEM, PBS, and 0.25% trypsin were from Corning. DMEM with no phenol red/glucose/ glutamine, Opti-MEM, GlutaMAX, 0.1mg/mL poly-D-lysine, and penicillin-streptomycin were from ThermoFisher Scientific. FBS was from GeminiBio. Lipofectamine 2000 was from Invitrogen. Rabbit anti-FLAG (D6W5B), Mouse anti-HA (6E2), Mouse anti-Beta Tubulin (D3U1W), Rabbit anti-GAPDH (14C10), and HRP-conjugated Rabbit anti-GAPDH (D16H11) XP antibodies were from Cell Signaling Technology. IRDye 800-conjugated Donkey Anti-Rabbit and IRDye 680-conjugated Goat Anti-Mouse secondary antibodies were from LI-COR Biosciences. To generate 647-conjugated Mouse anti-HA antibody, the antibody was purified from hybridoma supernatant and chemically conjugated to Alexa Fluor 647 NHS Ester from ThermoFisher Scientific. The Dual-Luciferase Reporter Assay System kit was from Promega. Coelenterazine 400a was from Nanolight Technology (Cat# 340).

### Plasmid DNA Constructs

Detailed construct features and calculated molecular weights for proteins used in this study are listed in Table S1.

All proteins were assembled using human aGPCR and G protein sequences retrieved from the Uniprot database. Synthetic gene fragments (Integrated DNA Technologies) were inserted into vectors using Gibson assembly, and plasmid sequences were confirmed by DNA sequencing. Receptor constructs were subcloned into the pFuse-hIgG1-Fc2 vector (Invivogen) using EcoRI and BglII restriction sites. All receptor constructs included a 3’ stop codon preceding the hIgG1-Fc coding sequence. An “empty vector” construct (referred to as “EV”; see Table S1) was generated by placement of a stop codon immediately after the IL2-signal sequence of the pFuse-hIgG1-Fc2 vector. A single glycine-serine linker was inserted between any epitope tag and the coding sequence of the protein of interest. Point mutants were generated using a Q5 Site-Directed Mutagenesis Kit (New England Biolabs).

We generated luciferase reporter plasmids containing individual response elements, a minimal promoter (minP), and *luc2* firefly luciferase. These plasmids are identical to the pGL4.29[CRE/minP/*luc2P*], pGL4.30[NFAT-RE/minP/*luc2P*], pGL4.33[SRE/*luc2P*], pGL4.34[SRF-RE/*luc2P*] plasmids (Promega) except that *luc2* is used in place of *luc2P* in our system and that minP is included in our SRE and SRF-RE plasmids. To assemble these plasmids, synthetic gene fragments containing response elements were inserted into a pGL4.23[*luc2*/minP] vector obtained from Promega using KpnI and HindIII (for SRE, SRF-RE, and CRE) or KpnI and NcoI (for NFAT-RE, which included the minP in the synthetic gene fragment) restriction sites. The pRL-TK plasmid (Promega) was used to express *Renilla* luciferase.

Tricistronic plasmids encoding TRUPATH G protein biosensors were a gift from Dr. Justin English, University of Utah. These plasmids utilize identical protein sequences (for various Gα subunits, Gβ1, and Gγ2) to those initially reported.

### Cell Culture

Parental HEK293T and HEKΔ6 cells were cultured in DMEM supplemented with 10% FBS and 1% penicillin-streptomycin at 37°C in a 5% CO_2_, humidified incubator.

### Reporter Gene Assays

96-well clear bottom, white walled, tissue culture-treated plates were coated with 40 μL of 10 μg/mL aqueous poly-D-lysine solution for 15 minutes at room temperature followed by aspiration to remove excess solution. Cells were seeded at a density of 3 x 10^5^ per well in 100 μL complete media (DMEM/10% FBS/1% penicillin-streptomycin) and incubated overnight (16-20 hours). Cells in 50 μL fresh complete media were transfected by adding 20 μL of a solution containing plasmid DNA and Lipofectamine 2000 at a ratio of 3 μL per 1 μg DNA in Opti-MEM.

For the CRE and NFAT-RE assays, each well was transfected with 50 ng firefly reporter plasmid, 1 ng pRL-TK plasmid, and 10 ng receptor plasmid. For the SRE and SRF-RE assays, each well was transfected with 30 ng firefly reporter plasmid, 0.6 ng pRL-TK plasmid, and 30 ng receptor plasmid. For receptor DNA titration experiments, total DNA was balanced with empty vector (see Table S1).

After 4-6 hours, the transfection media was replaced with 100 μL complete media and cells incubated overnight (16-20 hours). Media was then replaced with 100 μL complete media (CRE and NFAT-RE) or serum-free DMEM supplemented with 1% penicillin-streptomycin (SRE and SRF-RE), and cells were incubated for 6-8 hours. Cells were then washed with 100 μL PBS and lysed by addition of 20 μL 1 x Passive Lysis Buffer (Promega) to each well and nutated for 15 minutes at room temperature. Lysates were pipetted up and down, and the assay plate was spun at 400 x g for 60 seconds to remove air bubbles. The ratio of firefly to *Renilla* luciferase activity was measured using automated reagent injection with a GloMax luminometer system (Promega), in which 50 μL LAR II and 25 μL Stop & Glo reagent (Promega) were added to each well.

### G Protein Biosensor Assays

Cells were seeded and transfected using identical methods to those used for reporter gene assays. Each well was transfected with 80 ng of biosensor plasmid and 20 ng of receptor plasmid. After 4-6 hours, the transfection media was replaced with 100 μL complete media and cells incubated overnight. After 24 hours, media was replaced with 100 μL DMEM with no phenol red, glucose, or glutamine supplemented with 2% GlutaMAX and 1% penicillin-streptomycin. 14-20 hours later, media was replaced with Hank’s Balanced Salt Solution with no phenol red, calcium, or magnesium and supplemented with 20 mM HEPES pH 7.4 and 5 μM coelenterazine 400a from NanoLight Technology (Cat# 340-5). An opaque white vinyl sticker was placed on the bottom of the plate and BRET ratios were measured 2-8 minutes later using a GloMax luminometer system. RLucII luminescence was measured using ET405/40x and ET510lp filters (Chroma Technology Corp), for donor (*Renilla*) and acceptor (GFP) readouts, respectively.

### Cell Surface Staining and Flow Cytometry

Quantification of cell surface staining by flow cytometry was performed using cells plated in non-coated 96-well plates at a density of 3 x 10^5^ per well in 100 μL complete media and incubated overnight (16-20 hours). Then 50 μL media was removed from each well, and cells were transfected by adding 20 μL of a solution containing 60 ng plasmid DNA and Lipofectamine 2000 at a ratio of 3 μL per 1 μg DNA in Opti-MEM. After 4-6 hours, 30 μL of fresh complete media was added to each well, and cells were incubated overnight (18-20 hours). 24 hours post-transfection, cells were resuspended and transferred to a 96-well V-bottom plate (Corning). Cells were pelleted at 500 x g for 3 minutes, washed once with HEPES buffered saline (HBS), pH 7.4, and then rocked at 4° C in 100 μL of HBS supplemented with 0.1% bovine serum albumin (HBS/BSA) and 647-conjugated mouse anti-HA antibody at 1 μg/mL. Cells were rocked at 4° C for 20 minutes, washed twice with HBS/BSA, and analyzed using a Beckman Coulter Cytoflex flow cytometer. Data were analyzed with CytExpert software. Single cells were gated using forward and side scattering, and α-HA surface staining was quantified using the MFI in the APC channel.

### Western Blot Analysis

Cells for western blots were plated on poly-D-lysine-treated 24-well plates and transfected in parallel with cells for reporter assays, using the same transfection mixtures (which included both reporter plasmids and receptor and/or secTEV plasmids), and treated identically. Cell density, media volumes, and DNA loads were increased five-fold relative to the amounts used in 96-well format reporter assays to account for the larger well size. At the time of harvest, protein samples for blots were reduced and denatured in Laemmli buffer, loaded into 4 –20% Tris-Glycine gels (Bio-Rad) for SDS-PAGE, and transferred onto nitrocellulose membranes (Cytiva). Blots were blocked with 5% milk in TBST and incubated with primary antibodies overnight at 4°C. Simultaneous analysis of proteins of interest and loading controls were carried out for each blot through the use of two imaging channels on a LI-COR Odyssey CLx Fluorescent Imaging System. Blots were incubated with secondary antibodies for 30-60 minutes prior to imaging. For blots simultaneously detecting FLAG, HA, and GAPDH (Fig. S4), FLAG and HA were detected using the LI-COR system and HRP-conjugated GAPDH primary antibody was incubated with blots for 2 hours at room temperature and detected with Western Lightning Plus-ECL (PerkinElmer) using a Bio-Rad Chemidoc System.

### Bioinformatics and Structural Model Generation

To generate a homology model of the ADGRE5 CTF in the active signaling state (Fig. S3C), the sequences of human ADGRE5 residues 531-835 and human Gα_13_ residues 1-377 were submitted to the ColabFold software server^40^, which predicted a structure of the TA-agonized CTF in complex with Gα_13_. ColabFold implementation of AlphaFold installed on SBGrid was used to make structural predictions of all 32 human aGPCR CTF proteins^39,40,57^. All 5 models were run for 5 recycles with version 1.3.0, without templates and without amber relaxation. Multiple sequence alignments were generated using MMseqs2^58^. Models were ranked by the Predicted Template Modeling Score (pTM) and the top ranked model for each human adhesion GPCR was used for further analysis.

### Heat Map Analysis and Scaling of Measured Responses

TA-dependent genetic reporter scores (for heat map and scatter plots) were quantified as Log1P-transformed differences in signal given by MBP-CTF and MBP-CTFΔΤΑ proteins. Values were min-max scaled from zero to one within each assay. G protein biosensor scores were quantified as the ΔBRET value, linearly scaled from zero to the maximum value given in each respective biosensor for heat maps or from zero to one for scatter plots. Negative ΔBRET values were represented as zero in both metrics. Signaling score (in scatter plots) is the largest score amongst all reporter gene scores and G protein biosensor scores. TA-dependent surface depletion was quantified as the percentage decrease in surface staining signal given by MBP-CTF proteins when compared to the cognate MBP-CTFΔΤΑ proteins. For scatter plots, this was scaled from zero (0% depletion) to one (100% depletion).

---

**Fig. S1.**
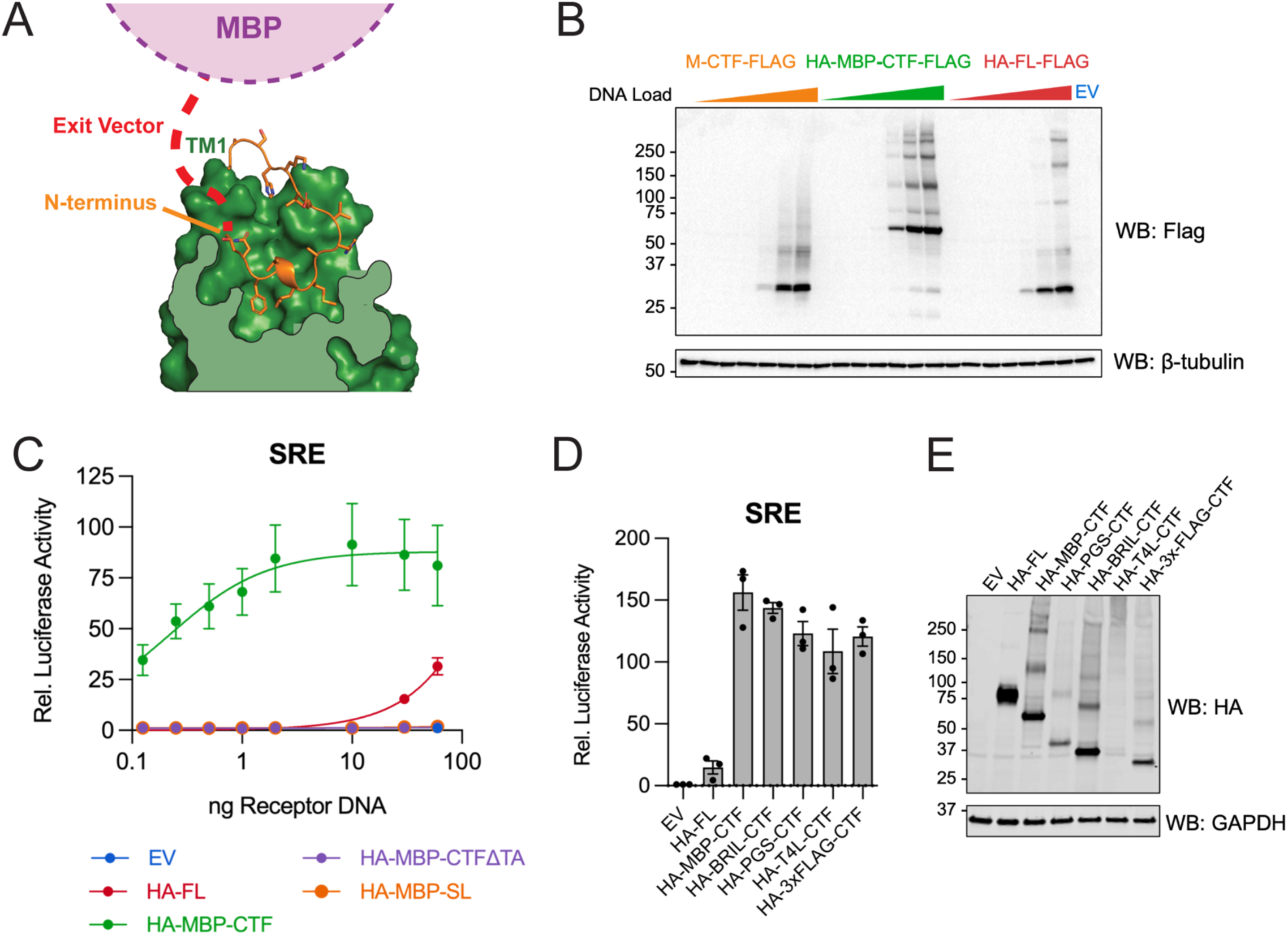
–Signaling properties of CTF fusion proteins. (A) Binding pose of the TA peptide within the 7TM domain of ADGRG1 (PDB ID: 7SF8)^34^. The presence of a potential exit vector path provides a physical explanation for the signaling capacity of CTF molecules with N-terminal fusions. (B) α-FLAG western blot analysis of protein abundance for molecules evaluated in the dose-response analysis of signaling in the SRE assay (Fig. 1D). Individual lanes correspond to DNA amounts used, with β-tubulin as a protein loading control. (C) Comparison of the indicated ADGRE5-derived proteins in the SRE assay as a function of the amount of transfected DNA. (D) SRE signaling activity for a panel of different N-terminal sequences fused to the ADGRE5 CTF. (E) Western blot detection of N-terminal HA epitope tags for fusion proteins evaluated in (D), with GADPH as a protein loading control. For panels C and D, all data points are normalized to the mean value for empty vector and presented as mean ± SEM of three independent biological replicates (n=3). In panel C, dose-response curves were fit to each titration curve with nonlinear regression.

**Fig. S2.**
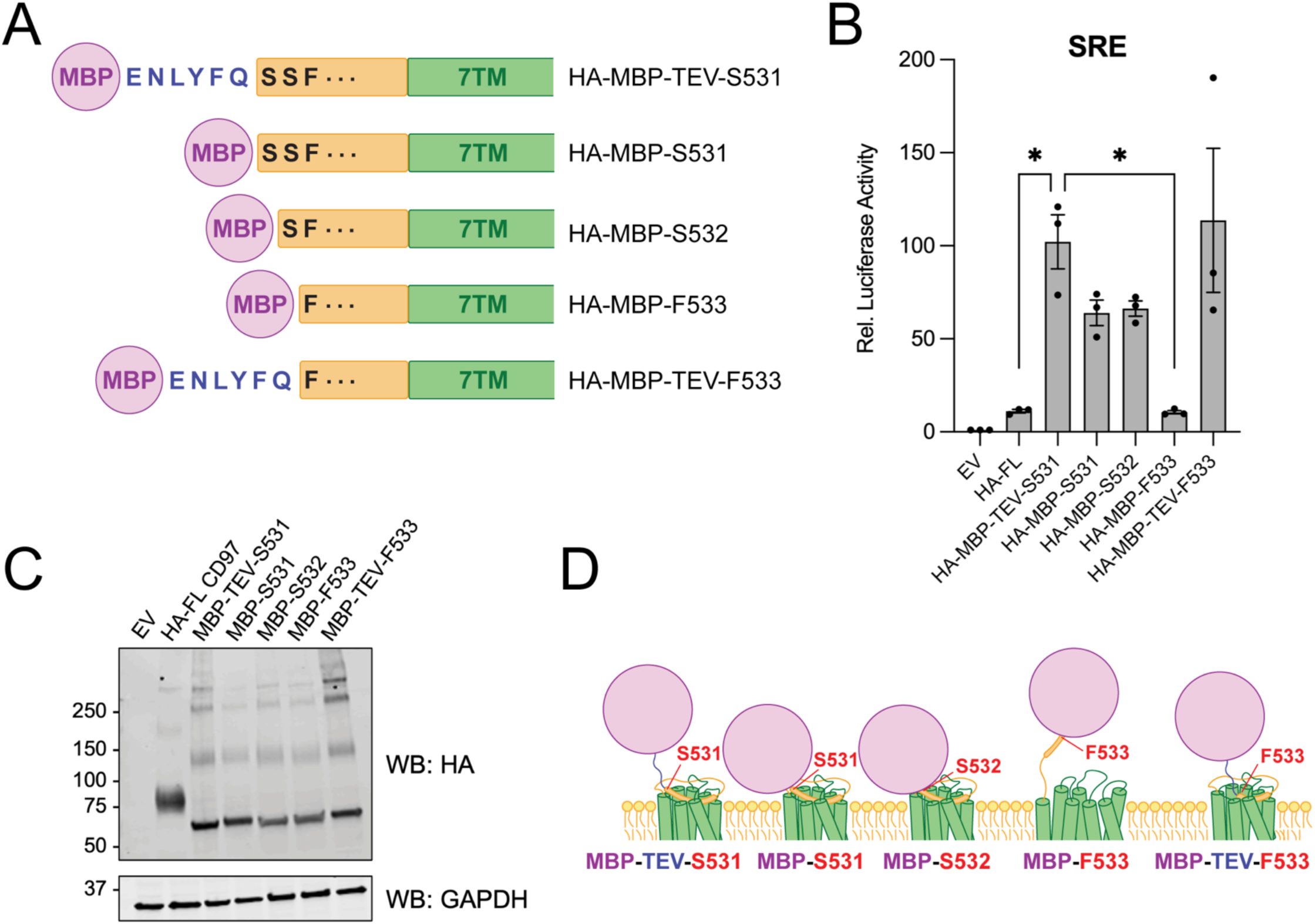
– Identification of the closest MBP fusion tolerable for autonomous aGPCR signaling in ADGRE5. (A) Schematic diagrams depicting constructs in which MBP is fused at decreasing distances from the TA. (B) Signaling response of MBP fusion constructs shown in (A). Data are normalized to the mean value for empty vector and presented as mean ± SEM of three independent biological replicates (n=3). One-way ANOVA was used with Tukey’s multiple-comparison post-hoc test to compare conditions (* P<0.05). (C) Western blot detection of N-terminal HA epitope tags for proteins evaluated in (B). Blotting for GAPDH was used to control for protein loading. (D) Model illustrating the variation in steric interference depending on the fusion position of MBP.

**Fig. S3.**
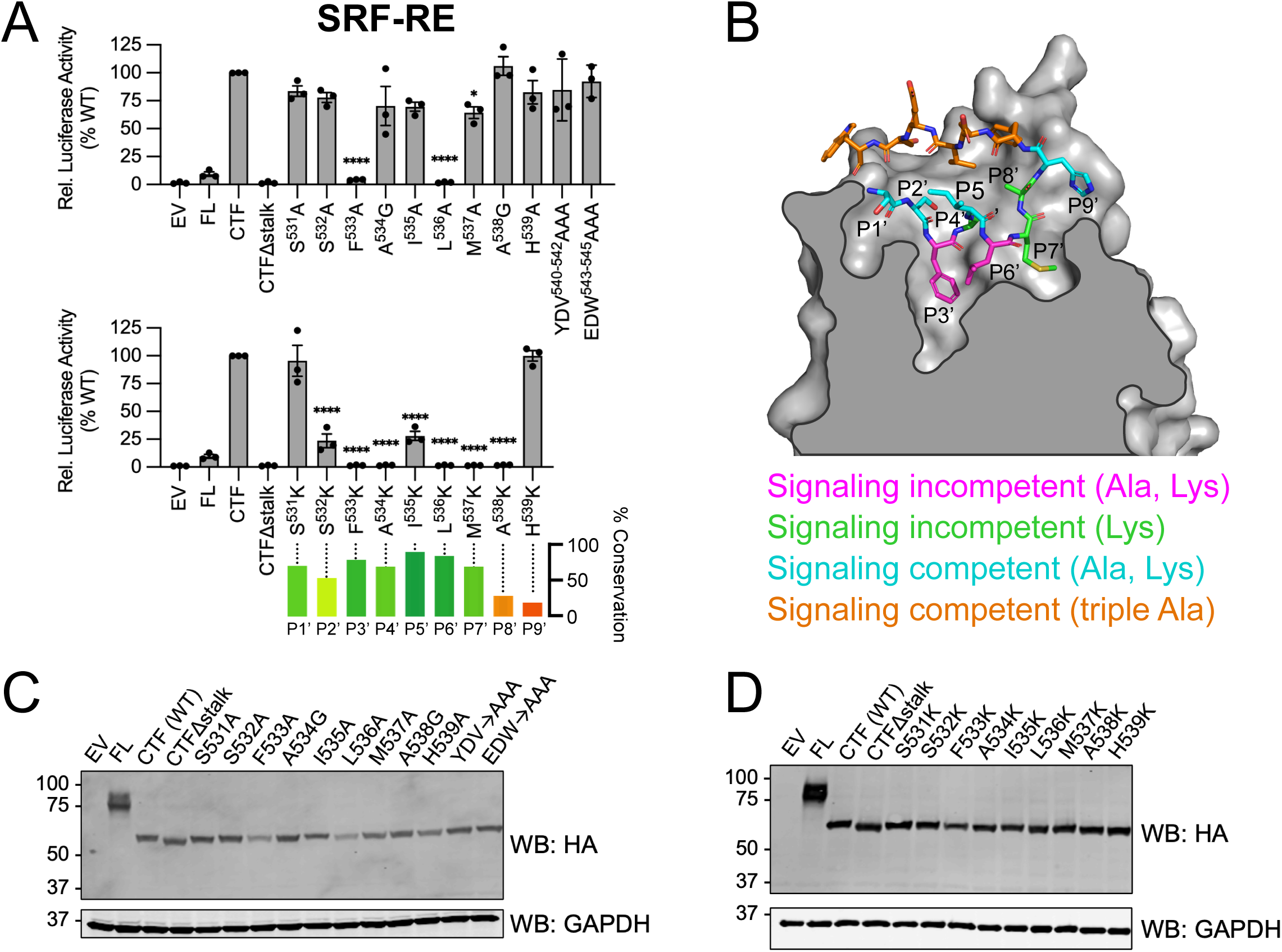
– Scanning mutagenesis evaluation of the ADGRE5 TA sequence in the MBP-CTF scaffold. (A) SRE signaling data for scanning alanine (or glycine for A534 and A538) and lysine mutagenesis of the stalk sequence in ADGRE5. All mutations were made in the context of parental MBP-CTF protein. Data are normalized as a percentage of the mean value for WT and presented as mean ± SEM of three independent biological replicates (n=3). Sequence conservation is shown below and based on alignment of the 33 human aGPCRs. One-way ANOVA was used with Tukey’s multiple-comparison post-hoc test to compare each mutant to wild-type protein (CTF) (* P<0.05, **** P<0.0001). (B) ColabFold-generated structural prediction of the TA-bound 7TM domain of ADGRE5 in complex with Gα_13_^40^. The orthosteric pocket on the extracellular face of the receptor is depicted, with the bound stalk sequence colored according to mutagenesis results. (C,D) Western blot detection of N-terminal HA epitope tags for constructs evaluated in (A). Blotting for GAPDH was used to control for protein loading.

**Fig. S4.**
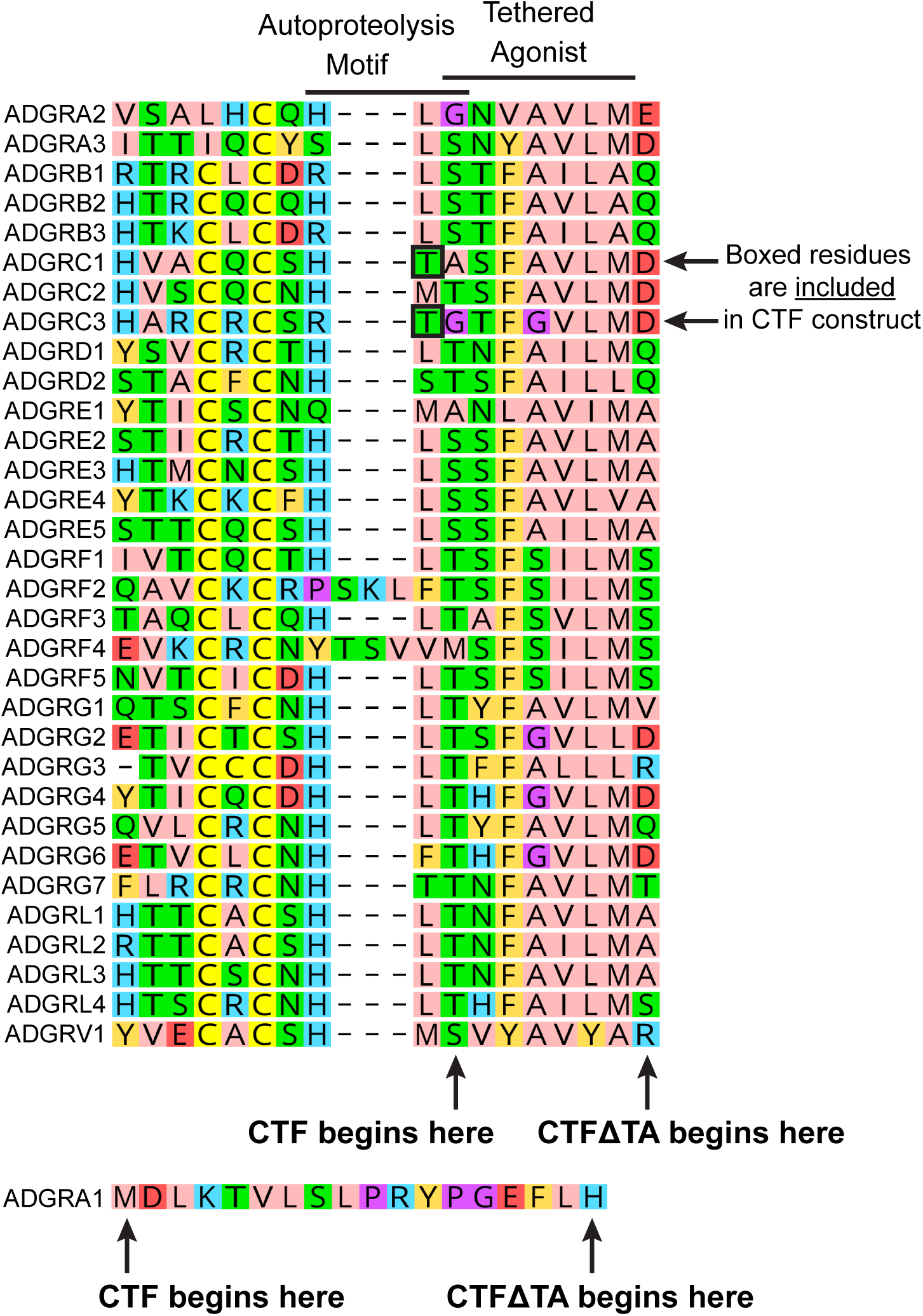
– Design of CTF and CTFΔTA protein constructs. A multiple sequence alignment of 32 human GAIN domains was used to identify the first residue of each CTF and CTFΔTA protein. The P1’ residue of each TA is the first residue of each CTF protein, except for ADGRC1 and ADGRC3, which start at the residue preceding P1’ because each possesses a nucleophilic threonine residue immediately after the basic residue of the autoproteolysis motif. The CTFΔTA proteins begin with the residue immediately following the P7’ TA position, as indicated. For ADGRA1, which lacks a GAIN domain, the CTF construct consists of the full-length receptor and the CTFΔTA construct lacks the 17 N-terminal residues prior to TM1.

**Fig. S5.**
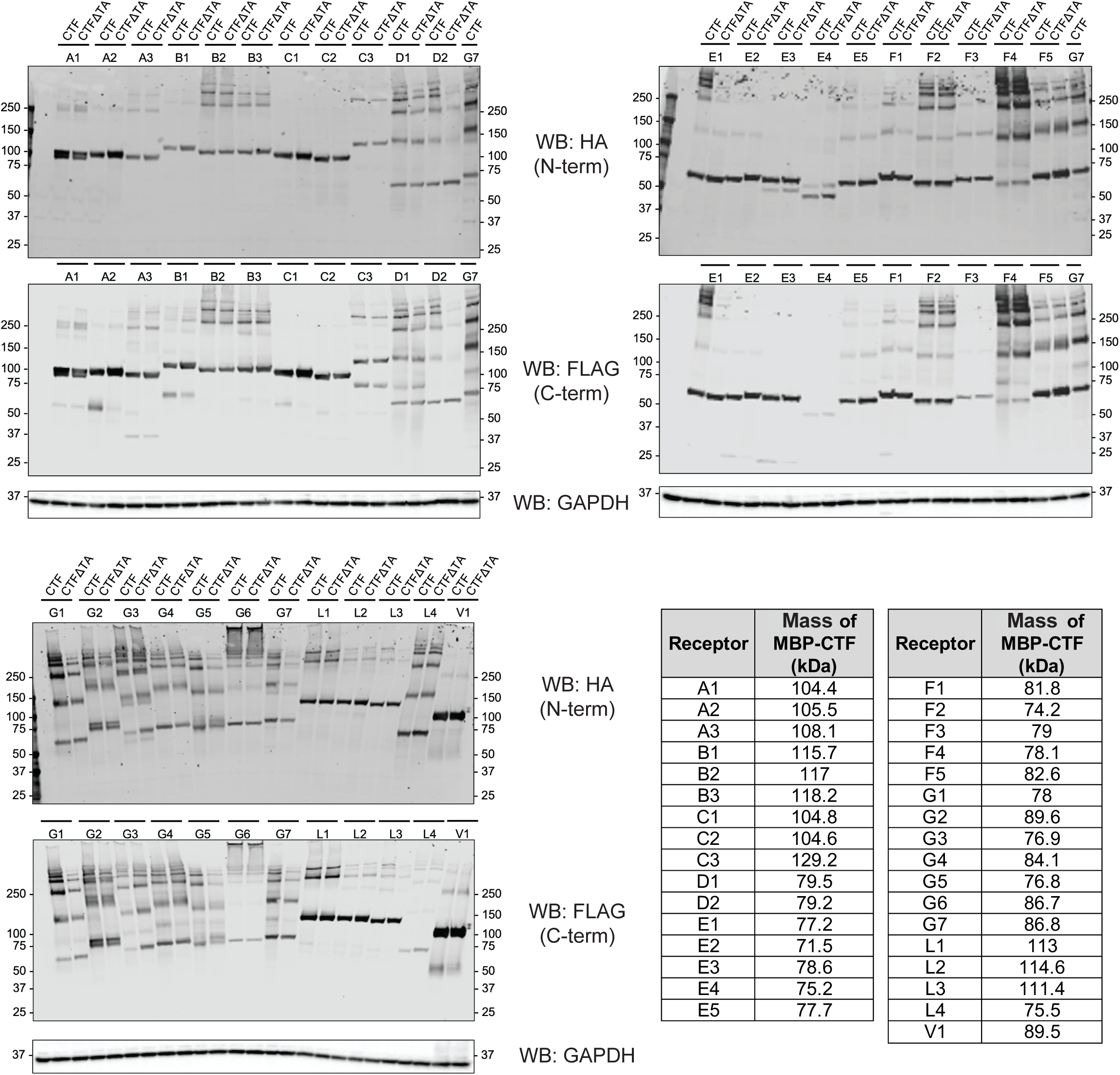
– Expression of human aGPCR proteins evaluated in this study. Western blot detection of N-terminal HA and C-terminal FLAG epitope tags to evaluate total protein expression for MBP-CTF and MBP-CTFΔΤΑ variants of all 33 human aGPCRs (Note: ADGRE4 is a predicted pseudogene and no mRNA transcripts encoding the 7TM domain have been detected in humans). Blotting for GAPDH was used to control for protein loading.

**Fig. S6.**
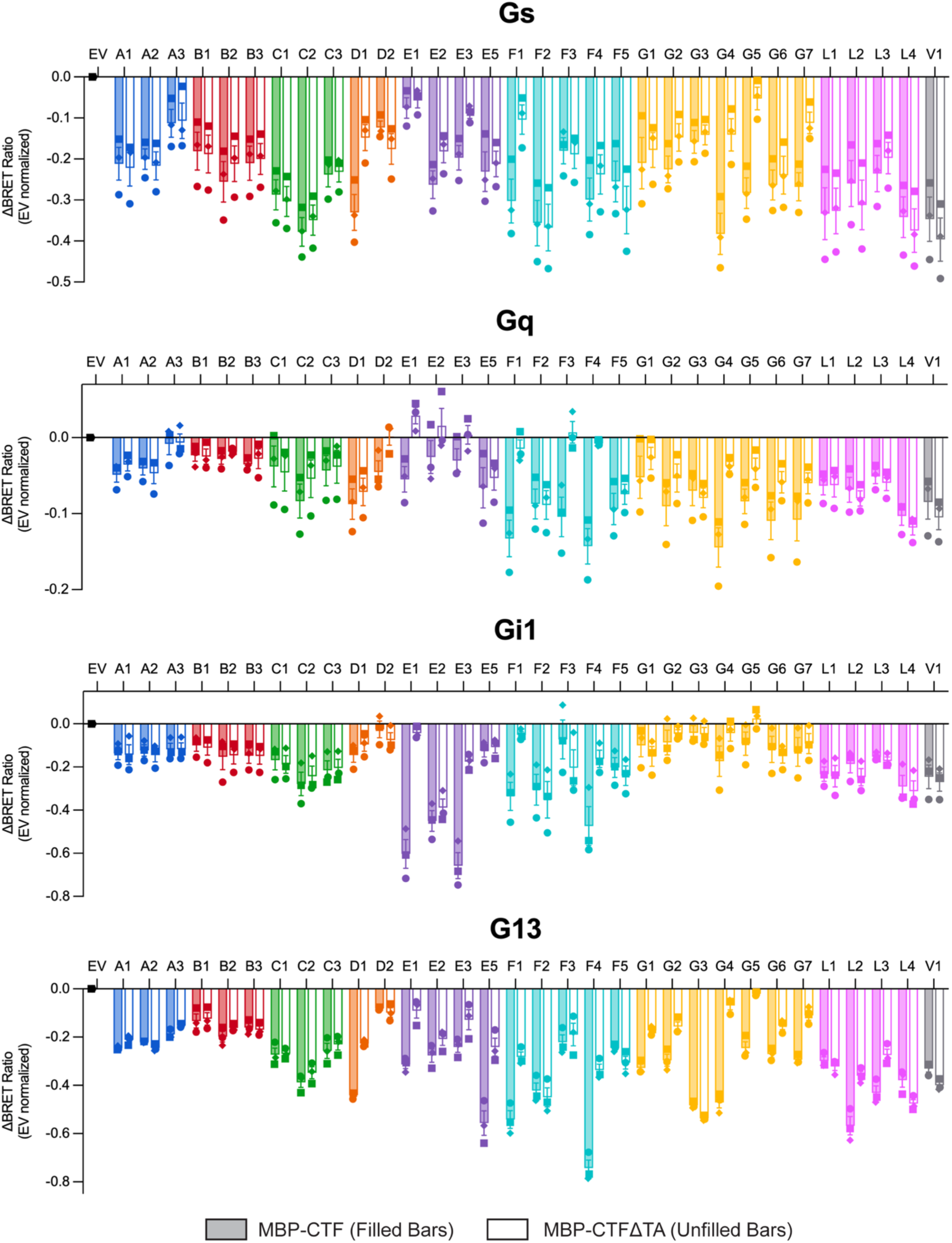
– Empty vector-normalized resting BRET ratios for all proteins tested in four G protein biosensor assays. Relative BRET ratios given by the human MBP-CTF (filled bars) and MBP-CTFΔTA (unfilled bars) proteins across the four G protein biosensors tested. For each protein, ΔBRET values are normalized to the resting BRET ratio established by cells transfected with empty vector. Data are presented as mean ± SEM of three independent biological replicates (n=3). Individual biological replicates are represented by circle, square, and diamond-shaped data points.

**Fig. S7.**
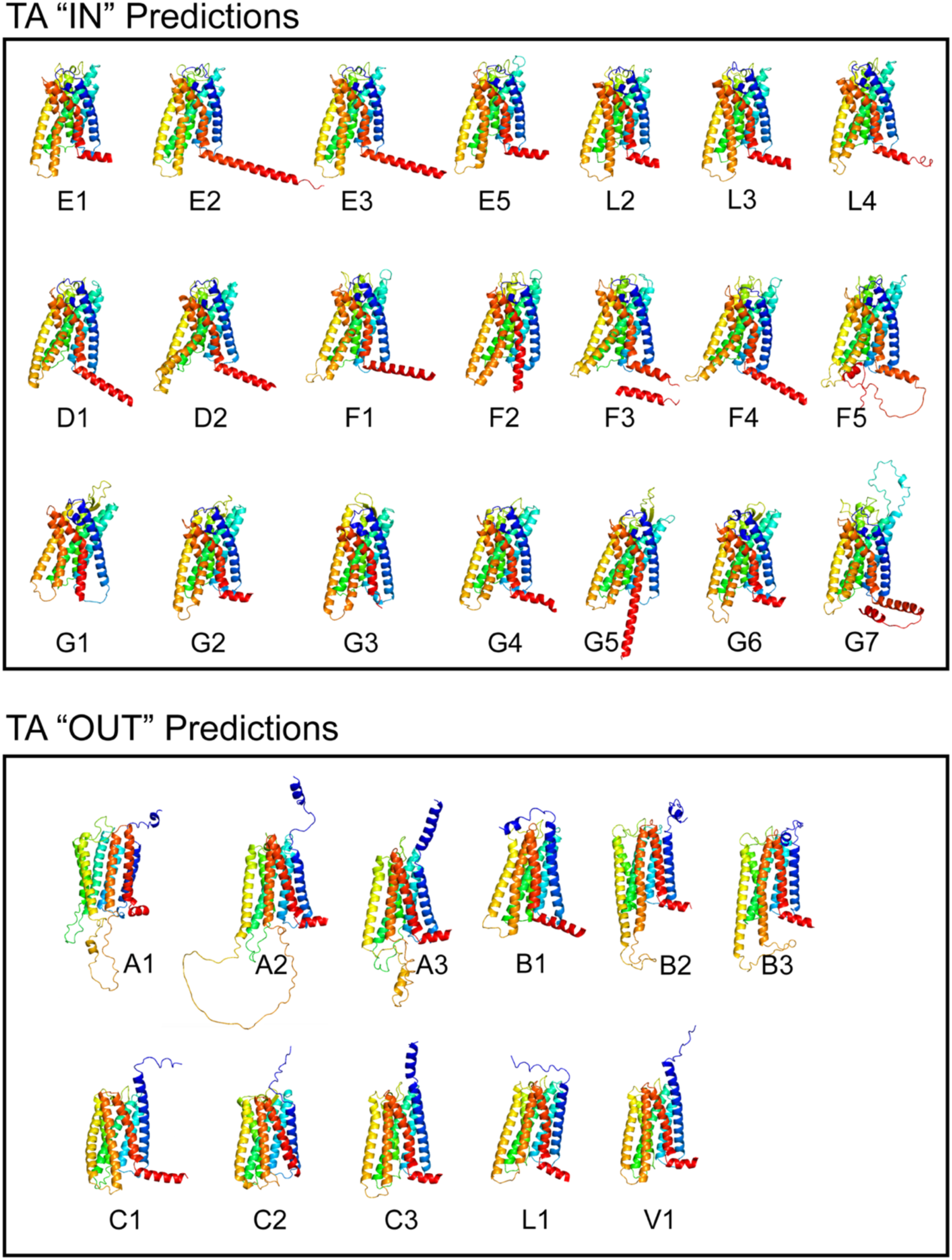
– ColabFold structural predictions for the human CTF proteins. Structural predictions for each CTF protein sequence from ColabFold. Models were manually sorted into two clusters based on the position of the TA peptide (in or out of the orthosteric pocket of the 7TM domain) and the presence of the kink in TM6, which is colored orange.

**Fig. S8.**
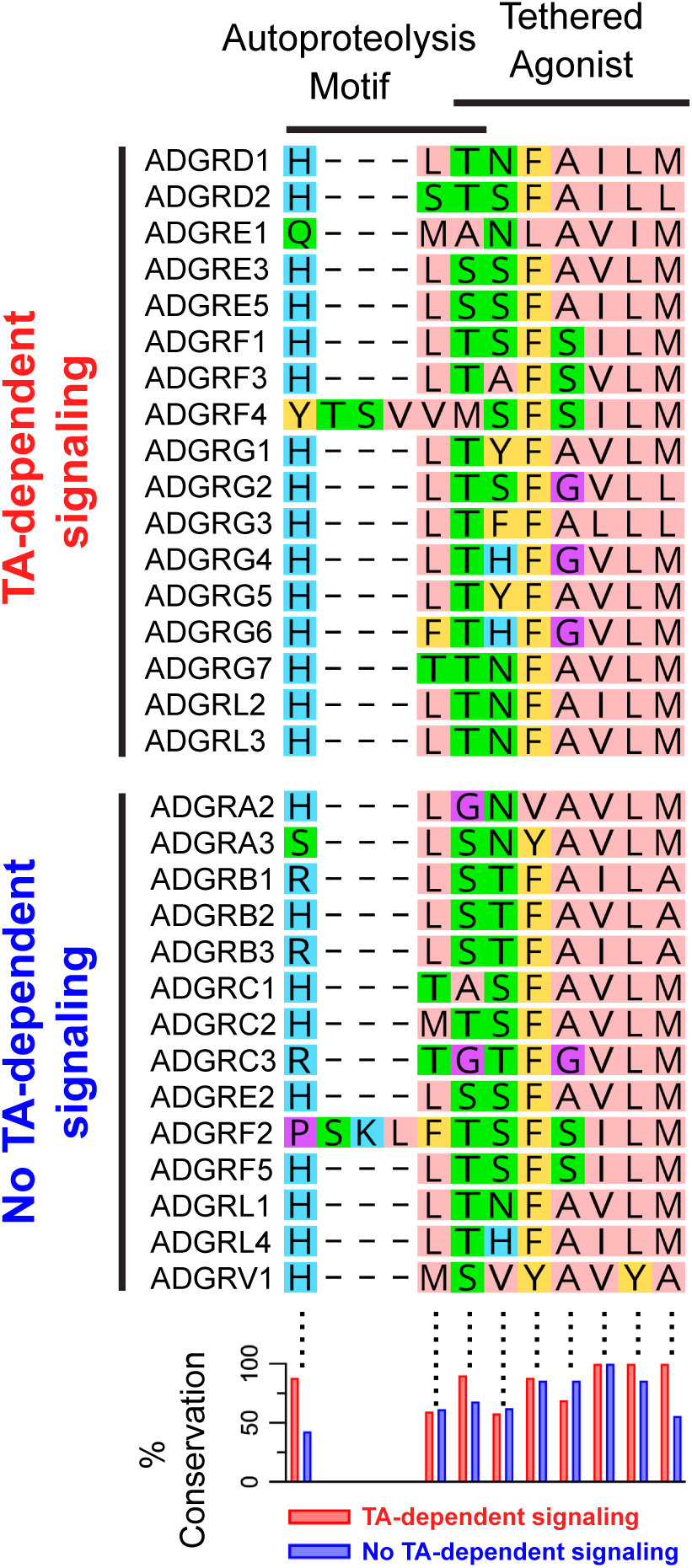
– Conservation of the TA sequence and autoproteolysis motif among aGPCRs. Multiple sequence alignment of aGPCRs around the autoproteolysis motif and the TA, with aGPCRs separated based on TA-signaling competence in our MBP fusion system. The amount of conservation as a function of position is plotted below the sequence alignment.

**Fig. S9.**
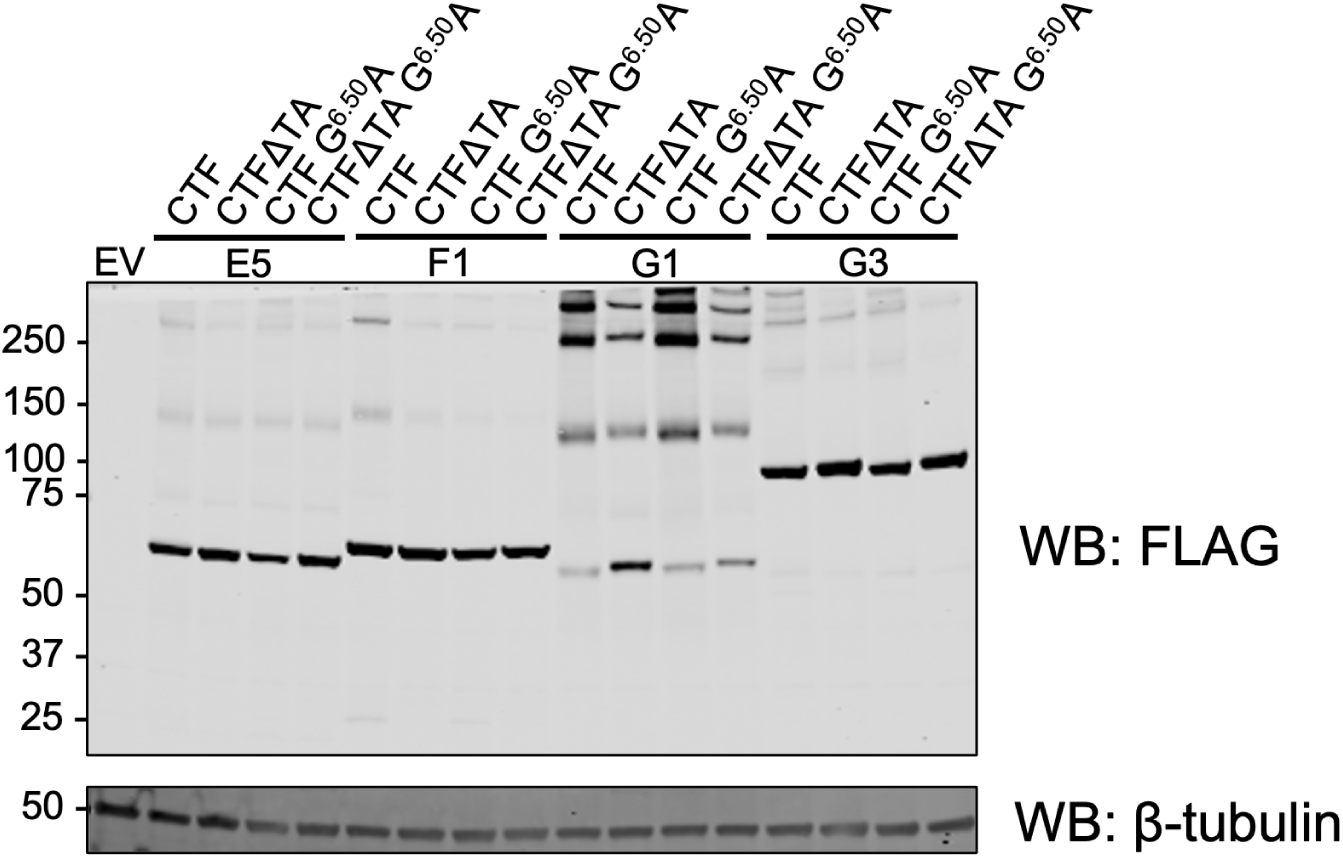
– Expression of G^6.50^A mutants. Western blot detection of C-terminal FLAG epitope tags for constructs evaluated in Fig. 6B. Blotting for β-tubulin was used to control for protein loading.

**Fig. S10.**
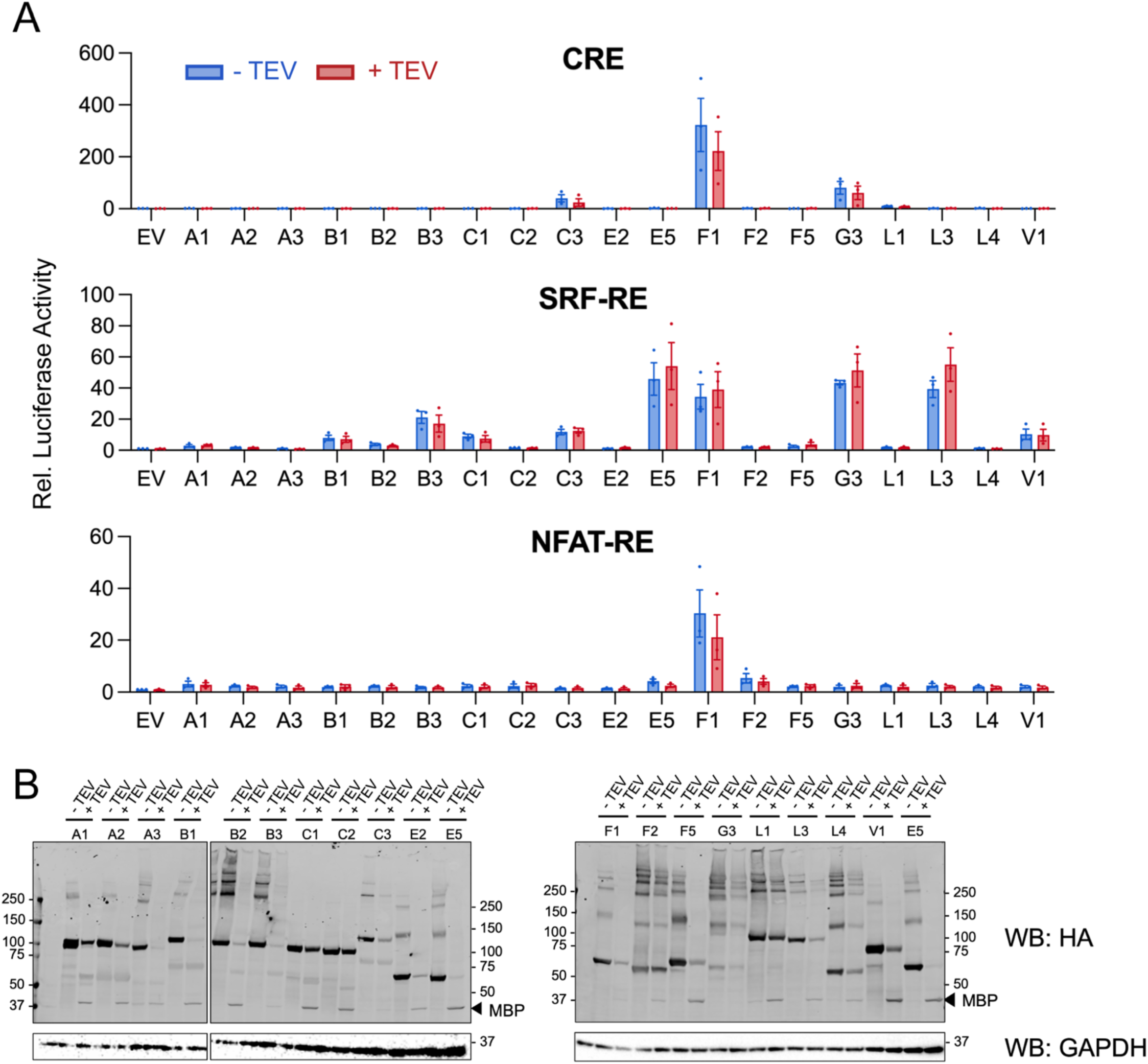
– Effects of TEV-mediated MBP removal on signaling activity of CTF proteins. (A) CRE, SRF-RE, and NFAT-RE reporter gene assays for cells co-expressing various MBP-CTF proteins with (red) and without (blue) a TEV protease variant engineered for activity in the secretory pathway (secTEV). Data are normalized to the mean value for empty vector without secTEV expression and are presented as mean ± SEM of three independent biological replicates (n=3). (B) Western Blot detection of N-terminal HA epitope tags on MBP-CTF proteins co-expressed with and without secTEV protease. Blotting for GAPDH was used to control for protein loading.

