## Supplemental Table 1 for "Heterogeneity of Tethered Agonist Signaling in Adhesion G Protein-Coupled Receptors"

| Name | Residues Included | Vector | Description | Calculated MW (if blotted) | Source |
|---|---|---|---|---|---|
| Empty Vector (EV) | | pFuse-hIgG1-Fc2 | Empty pFuse-hIgG1-Fc2 plasmid with a stop codon inserted between the IL2 signal peptide and the multiple cloning site | | |
| ADGRE5 M-CTF-FLAG | 531-835 | pFuse-hIgG1-Fc1 | Methionine residue followed by a ADGRE5 construct N-terminally truncated at the autoproteolytic cleavage site (ADGRE5 CTF) and a FLAG tag | 35.5 kDa | |
| ADGRE5 M-CTFΔstalk-FLAG | 546-835 | pFuse-hIgG1-Fc1 | Methionine residue followed by a ADGRE5 construct N-terminally truncated at the start of TM1 (ADGRE5 CTFΔstalk) and a FLAG tag | 33.8 kDa | |
| ADGRE5 M-CTFΔTA-FLAG | 538-835 | pFuse-hIgG1-Fc1 | Methionine residue followed by a ADGRE5 construct N-terminally truncated at the P8' residue (ADGRE5 CTFΔTA) and a FLAG tag | 34.8 kDa | |
| HA-MBP-ADGRE5 CTF-FLAG | 531-835 | pFuse-hIgG1-Fc2 | IL2 signal peptide followed by an HA tag, maltose binding protein, TEV cleavage sequence residues P6-P1, the ADGRE5 CTF, and a FLAG tag | 77.7 kDa | |
| HA-MBP-ADGRE5 SL-FLAG | 546-835 | pFuse-hIgG1-Fc2 | IL2 signal peptide followed by an HA tag, maltose binding protein, TEV cleavage sequence residues P6-P1, ADGRE5 SL, and a FLAG tag | 76.0 kDa | |
| HA-ADGRE5 FL-FLAG | 21-835 | pFuse-hIgG1-Fc2 | IL2 signal peptide followed by an HA tag, full-length ADGRE5 isoform 1, and a FLAG tag | 92.1 kDa (unprocessed); 56.7 kDa (NTF); 35.4 kDa (CTF) | |
| SRE/MinP/*luc2* | | pGL4.23 | Luciferase reporter vector using the serum response element, a minimal promoter, and luc2 firefly luciferase (modified from Promega pGL4.33 vector). | | Ref. 44 |
| SRF-RE/MinP/*luc2* | | pGL4.23 | Luciferase reporter vector using the serum response factor response element, a minimal promoter, and luc2 firefly luciferase (modified from Promega pGL4.34 vector). | | Ref. 44 |
| NFAT-RE/MinP/*luc2* | | pGL4.23 | Luciferase reporter vector using the nuclear factor of activated T cells response element, a minimal promoter, and luc2 firefly luciferase (modified from Promega pGL4.30 vector). | | Ref. 44 |
| CRE/MinP/*luc2* | | pGL4.23 | Luciferase reporter vector using the cAMP response element, a minimal promoter, and luc2 firefly luciferase (modified from Promega pGL4.29 vector). | | Ref. 44 |
| HA-MBP-ADGRA1 CTF-FLAG | 1-560 | pFuse-hIgG1-Fc2 | IL2 signal peptide followed by an HA tag, maltose binding protein, TEV cleavage sequence residues P6-P1, the relevant CTF sequence, and a FLAG tag | 104.4 kDa | |
| HA-MBP-ADGRA1 CTFΔTA-FLAG | 18-560 | pFuse-hIgG1-Fc2 | IL2 signal peptide followed by an HA tag, maltose binding protein, TEV cleavage sequence residues P6-P1, the relevant CTFΔTA sequence, and a FLAG tag | 102.4 kDa | |
| HA-MBP-ADGRA2 CTF-FLAG | 747-1338 | pFuse-hIgG1-Fc2 | IL2 signal peptide followed by an HA tag, maltose binding protein, TEV cleavage sequence residues P6-P1, the relevant CTF sequence, and a FLAG tag | 105.5kDa | |
| HA-MBP-ADGRA2 CTFΔTA-FLAG | 754-1338 | pFuse-hIgG1-Fc2 | IL2 signal peptide followed by an HA tag, maltose binding protein, TEV cleavage sequence residues P6-P1, the relevant CTFΔTA sequence, and a FLAG tag | 104.8kDa | |
| HA-MBP-ADGRA3 CTF-FLAG | 738-1321 | pFuse-hIgG1-Fc2 | IL2 signal peptide followed by an HA tag, maltose binding protein, TEV cleavage sequence residues P6-P1, the relevant CTF sequence, and a FLAG tag | 108.1kDa | |
| HA-MBP-ADGRA3 CTFΔTA-FLAG | 745-1321 | pFuse-hIgG1-Fc2 | IL2 signal peptide followed by an HA tag, maltose binding protein, TEV cleavage sequence residues P6-P1, the relevant CTFΔTA sequence, and a FLAG tag | 107.4kDa | |
| HA-MBP-ADGRB1 CTF-FLAG | 927-1584 | pFuse-hIgG1-Fc2 | IL2 signal peptide followed by an HA tag, maltose binding protein, TEV cleavage sequence residues P6-P1, the relevant CTF sequence, and a FLAG tag | 115.7kDa | |
| HA-MBP-ADGRB1 CTFΔTA-FLAG | 934-1584 | pFuse-hIgG1-Fc2 | IL2 signal peptide followed by an HA tag, maltose binding protein, TEV cleavage sequence residues P6-P1, the relevant CTFΔTA sequence, and a FLAG tag | 115kDa | |
| HA-MBP-ADGRB2 CTF-FLAG | 912-1585 | pFuse-hIgG1-Fc2 | IL2 signal peptide followed by an HA tag, maltose binding protein, TEV cleavage sequence residues P6-P1, the relevant CTF sequence, and a FLAG tag | 117kDa | |
| HA-MBP-ADGRB2 CTFΔTA-FLAG | 919-1585 | pFuse-hIgG1-Fc2 | IL2 signal peptide followed by an HA tag, maltose binding protein, TEV cleavage sequence residues P6-P1, the relevant CTFΔTA sequence, and a FLAG tag | 116.3kDa | |
| HA-MBP-ADGRB3 CTF-FLAG | 857-1522 | pFuse-hIgG1-Fc2 | IL2 signal peptide followed by an HA tag, maltose binding protein, TEV cleavage sequence residues P6-P1, the relevant CTF sequence, and a FLAG tag | 118.2kDa | |
| HA-MBP-ADGRB3 CTFΔTA-FLAG | 864-1522 | pFuse-hIgG1-Fc2 | IL2 signal peptide followed by an HA tag, maltose binding protein, TEV cleavage sequence residues P6-P1, the relevant CTFΔTA sequence, and a FLAG tag | 117.5kDa | |
| HA-MBP-ADGRC1 CTF-FLAG | 2448-3014 | pFuse-hIgG1-Fc2 | IL2 signal peptide followed by an HA tag, maltose binding protein, TEV cleavage sequence residues P6-P1, the relevant CTF sequence, and a FLAG tag | 104.8kDa | |
| HA-MBP-ADGRC1 CTFΔTA-FLAG | 2456-3014 | pFuse-hIgG1-Fc2 | IL2 signal peptide followed by an HA tag, maltose binding protein, TEV cleavage sequence residues P6-P1, the relevant CTFΔTA sequence, and a FLAG tag | 104.1kDa | |
| HA-MBP-ADGRC2 CTF-FLAG | 2357-2923 | pFuse-hIgG1-Fc2 | IL2 signal peptide followed by an HA tag, maltose binding protein, TEV cleavage sequence residues P6-P1, the relevant CTF sequence, and a FLAG tag | 104.6kDa | |

| | | | | |
|---|---|---|---|---|
| HA-MBP-ADGRC2 CTFΔTA-FLAG | 2364-2923 | pFuse-hIgG1-Fc2 | IL2 signal peptide followed by an HA tag, maltose binding protein, TEV cleavage sequence residues P6-P1, the relevant CTFΔTA sequence, and a FLAG tag | 103.9kDa |
| HA-MBP-ADGRC3 CTF-FLAG | 2517-3312 | pFuse-hIgG1-Fc2 | IL2 signal peptide followed by an HA tag, maltose binding protein, TEV cleavage sequence residues P6-P1, the relevant CTF sequence, and a FLAG tag | 129.2kDa |
| HA-MBP-ADGRC3 CTFΔTA-FLAG | 2525-3312 | pFuse-hIgG1-Fc2 | IL2 signal peptide followed by an HA tag, maltose binding protein, TEV cleavage sequence residues P6-P1, the relevant CTFΔTA sequence, and a FLAG tag | 128.5kDa |
| HA-MBP-ADGRD1 CTF-FLAG | 545-874 | pFuse-hIgG1-Fc2 | IL2 signal peptide followed by an HA tag, maltose binding protein, TEV cleavage sequence residues P6-P1, the relevant CTF sequence, and a FLAG tag | 79.5kDa |
| HA-MBP-ADGRD1 CTFΔTA-FLAG | 552-874 | pFuse-hIgG1-Fc2 | IL2 signal peptide followed by an HA tag, maltose binding protein, TEV cleavage sequence residues P6-P1, the relevant CTFΔTA sequence, and a FLAG tag | 78.8kDa |
| HA-MBP-ADGRD2 CTF-FLAG | 637-963 | pFuse-hIgG1-Fc2 | IL2 signal peptide followed by an HA tag, maltose binding protein, TEV cleavage sequence residues P6-P1, the relevant CTF sequence, and a FLAG tag | 79.2kDa |
| HA-MBP-ADGRD2 CTFΔTA-FLAG | 644-963 | pFuse-hIgG1-Fc2 | IL2 signal peptide followed by an HA tag, maltose binding protein, TEV cleavage sequence residues P6-P1, the relevant CTFΔTA sequence, and a FLAG tag | 78.5kDa |
| HA-MBP-ADGRE1 CTF-FLAG | 585-886 | pFuse-hIgG1-Fc2 | IL2 signal peptide followed by an HA tag, maltose binding protein, TEV cleavage sequence residues P6-P1, the relevant CTF sequence, and a FLAG tag | 77.2kDa |
| HA-MBP-ADGRE1 CTFΔTA-FLAG | 592-886 | pFuse-hIgG1-Fc2 | IL2 signal peptide followed by an HA tag, maltose binding protein, TEV cleavage sequence residues P6-P1, the relevant CTFΔTA sequence, and a FLAG tag | 76.5kDa |
| HA-MBP-ADGRE2 CTF-FLAG | 518-823 | pFuse-hIgG1-Fc2 | IL2 signal peptide followed by an HA tag, maltose binding protein, TEV cleavage sequence residues P6-P1, the relevant CTF sequence, and a FLAG tag | 71.5kDa |
| HA-MBP-ADGRE2 CTFΔTA-FLAG | 525-823 | pFuse-hIgG1-Fc2 | IL2 signal peptide followed by an HA tag, maltose binding protein, TEV cleavage sequence residues P6-P1, the relevant CTFΔTA sequence, and a FLAG tag | 70.8kDa |
| HA-MBP-ADGRE3 CTF-FLAG | 339-652 | pFuse-hIgG1-Fc2 | IL2 signal peptide followed by an HA tag, maltose binding protein, TEV cleavage sequence residues P6-P1, the relevant CTF sequence, and a FLAG tag | 78.6kDa |
| HA-MBP-ADGRE3 CTFΔTA-FLAG | 346-652 | pFuse-hIgG1-Fc2 | IL2 signal peptide followed by an HA tag, maltose binding protein, TEV cleavage sequence residues P6-P1, the relevant CTFΔTA sequence, and a FLAG tag | 77.9kDa |
| HA-MBP-ADGRE4 CTF-FLAG | 174-457 | pFuse-hIgG1-Fc2 | IL2 signal peptide followed by an HA tag, maltose binding protein, TEV cleavage sequence residues P6-P1, the relevant CTF sequence, and a FLAG tag | 75.2kDa |
| HA-MBP-ADGRE4 CTFΔTA-FLAG | 181-457 | pFuse-hIgG1-Fc2 | IL2 signal peptide followed by an HA tag, maltose binding protein, TEV cleavage sequence residues P6-P1, the relevant CTFΔTA sequence, and a FLAG tag | 74.5kDa |
| HA-MBP-ADGRE5 CTF-FLAG (also listed above on row 5) | 531-835 | pFuse-hIgG1-Fc2 | IL2 signal peptide followed by an HA tag, maltose binding protein, TEV cleavage sequence residues P6-P1, the relevant CTF sequence, and a FLAG tag | 77.7kDa |
| HA-MBP-ADGRE5 CTFΔTA-FLAG | 538-835 | pFuse-hIgG1-Fc2 | IL2 signal peptide followed by an HA tag, maltose binding protein, TEV cleavage sequence residues P6-P1, the relevant CTFΔTA sequence, and a FLAG tag | 77kDa |
| HA-MBP-ADGRF1 CTF-FLAG | 567-910 | pFuse-hIgG1-Fc2 | IL2 signal peptide followed by an HA tag, maltose binding protein, TEV cleavage sequence residues P6-P1, the relevant CTF sequence, and a FLAG tag | 81.8kDa |
| HA-MBP-ADGRF1 CTFΔTA-FLAG | 574-910 | pFuse-hIgG1-Fc2 | IL2 signal peptide followed by an HA tag, maltose binding protein, TEV cleavage sequence residues P6-P1, the relevant CTFΔTA sequence, and a FLAG tag | 81.1kDa |
| HA-MBP-ADGRF2 CTF-FLAG | 430-708 | pFuse-hIgG1-Fc2 | IL2 signal peptide followed by an HA tag, maltose binding protein, TEV cleavage sequence residues P6-P1, the relevant CTF sequence, and a FLAG tag | 74.2kDa |
| HA-MBP-ADGRF2 CTFΔTA-FLAG | 437-708 | pFuse-hIgG1-Fc2 | IL2 signal peptide followed by an HA tag, maltose binding protein, TEV cleavage sequence residues P6-P1, the relevant CTFΔTA sequence, and a FLAG tag | 73.5kDa |
| HA-MBP-ADGRF3 CTF-FLAG | 753-1079 | pFuse-hIgG1-Fc2 | IL2 signal peptide followed by an HA tag, maltose binding protein, TEV cleavage sequence residues P6-P1, the relevant CTF sequence, and a FLAG tag | 79kDa |
| HA-MBP-ADGRF3 CTFΔTA-FLAG | 760-1079 | pFuse-hIgG1-Fc2 | IL2 signal peptide followed by an HA tag, maltose binding protein, TEV cleavage sequence residues P6-P1, the relevant CTFΔTA sequence, and a FLAG tag | 78.3kDa |
| HA-MBP-ADGRF4 CTF-FLAG | 385-695 | pFuse-hIgG1-Fc2 | IL2 signal peptide followed by an HA tag, maltose binding protein, TEV cleavage sequence residues P6-P1, the relevant CTF sequence, and a FLAG tag | 78.1kDa |
| HA-MBP-ADGRF4 CTFΔTA-FLAG | 392-695 | pFuse-hIgG1-Fc2 | IL2 signal peptide followed by an HA tag, maltose binding protein, TEV cleavage sequence residues P6-P1, the relevant CTFΔTA sequence, and a FLAG tag | 77.4kDa |
| HA-MBP-ADGRF5 CTF-FLAG | 991-1346 | pFuse-hIgG1-Fc2 | IL2 signal peptide followed by an HA tag, maltose binding protein, TEV cleavage sequence residues P6-P1, the relevant CTF sequence, and a FLAG tag | 82.6kDa |
| HA-MBP-ADGRF5 CTFΔTA-FLAG | 998-1346 | pFuse-hIgG1-Fc2 | IL2 signal peptide followed by an HA tag, maltose binding protein, TEV cleavage sequence residues P6-P1, the relevant CTFΔTA sequence, and a FLAG tag | 81.9kDa |
| HA-MBP-ADGRG1 CTF-FLAG | 383-693 | pFuse-hIgG1-Fc2 | IL2 signal peptide followed by an HA tag, maltose binding protein, TEV cleavage sequence residues P6-P1, the relevant CTF sequence, and a FLAG tag | 78kDa |
| HA-MBP-ADGRG1 CTFΔTA-FLAG | 390-693 | pFuse-hIgG1-Fc2 | IL2 signal peptide followed by an HA tag, maltose binding protein, TEV cleavage sequence residues P6-P1, the relevant CTFΔTA sequence, and a FLAG tag | 77.3kDa |
| HA-MBP-ADGRG2 CTF-FLAG | 607-1017 | pFuse-hIgG1-Fc2 | IL2 signal peptide followed by an HA tag, maltose binding protein, TEV cleavage sequence residues P6-P1, the relevant CTF sequence, and a FLAG tag | 89.6kDa |

| | | | | | |
|---|---|---|---|---|---|
| HA-MBP-ADGRG2 CTFΔTA-FLAG | 614-1017 | pFuse-hIgG1-Fc2 | IL2 signal peptide followed by an HA tag, maltose binding protein, TEV cleavage sequence residues P6-P1, the relevant CTFΔTA sequence, and a FLAG tag | 88.9kDa | |
| HA-MBP-ADGRG3 CTF-FLAG | 250-549 | pFuse-hIgG1-Fc2 | IL2 signal peptide followed by an HA tag, maltose binding protein, TEV cleavage sequence residues P6-P1, the relevant CTF sequence, and a FLAG tag | 76.9kDa | |
| HA-MBP-ADGRG3 CTFΔTA-FLAG | 257-549 | pFuse-hIgG1-Fc2 | IL2 signal peptide followed by an HA tag, maltose binding protein, TEV cleavage sequence residues P6-P1, the relevant CTFΔTA sequence, and a FLAG tag | 76.2kDa | |
| HA-MBP-ADGRG4 CTF-FLAG | 2722-3080 | pFuse-hIgG1-Fc2 | IL2 signal peptide followed by an HA tag, maltose binding protein, TEV cleavage sequence residues P6-P1, the relevant CTF sequence, and a FLAG tag | 84.1kDa | |
| HA-MBP-ADGRG4 CTFΔTA-FLAG | 2729-3080 | pFuse-hIgG1-Fc2 | IL2 signal peptide followed by an HA tag, maltose binding protein, TEV cleavage sequence residues P6-P1, the relevant CTFΔTA sequence, and a FLAG tag | 83.4kDa | |
| HA-MBP-ADGRG5 CTF-FLAG | 227-528 | pFuse-hIgG1-Fc2 | IL2 signal peptide followed by an HA tag, maltose binding protein, TEV cleavage sequence residues P6-P1, the relevant CTF sequence, and a FLAG tag | 76.8kDa | |
| HA-MBP-ADGRG5 CTFΔTA-FLAG | 234-528 | pFuse-hIgG1-Fc2 | IL2 signal peptide followed by an HA tag, maltose binding protein, TEV cleavage sequence residues P6-P1, the relevant CTFΔTA sequence, and a FLAG tag | 76.1kDa | |
| HA-MBP-ADGRG6 CTF-FLAG | 841-1221 | pFuse-hIgG1-Fc2 | IL2 signal peptide followed by an HA tag, maltose binding protein, TEV cleavage sequence residues P6-P1, the relevant CTF sequence, and a FLAG tag | 86.7kDa | |
| HA-MBP-ADGRG6 CTFΔTA-FLAG | 848-1221 | pFuse-hIgG1-Fc2 | IL2 signal peptide followed by an HA tag, maltose binding protein, TEV cleavage sequence residues P6-P1, the relevant CTFΔTA sequence, and a FLAG tag | 86kDa | |
| HA-MBP-ADGRG7 CTF-FLAG | 416-797 | pFuse-hIgG1-Fc2 | IL2 signal peptide followed by an HA tag, maltose binding protein, TEV cleavage sequence residues P6-P1, the relevant CTF sequence, and a FLAG tag | 86.8kDa | |
| HA-MBP-ADGRG7 CTFΔTA-FLAG | 423-797 | pFuse-hIgG1-Fc2 | IL2 signal peptide followed by an HA tag, maltose binding protein, TEV cleavage sequence residues P6-P1, the relevant CTFΔTA sequence, and a FLAG tag | 86.1kDa | |
| HA-MBP-ADGRL1 CTF-FLAG | 839-1474 | pFuse-hIgG1-Fc2 | IL2 signal peptide followed by an HA tag, maltose binding protein, TEV cleavage sequence residues P6-P1, the relevant CTF sequence, and a FLAG tag | 113kDa | |
| HA-MBP-ADGRL1 CTFΔTA-FLAG | 846-1474 | pFuse-hIgG1-Fc2 | IL2 signal peptide followed by an HA tag, maltose binding protein, TEV cleavage sequence residues P6-P1, the relevant CTFΔTA sequence, and a FLAG tag | 112.3kDa | |
| HA-MBP-ADGRL2 CTF-FLAG | 825-1459 | pFuse-hIgG1-Fc2 | IL2 signal peptide followed by an HA tag, maltose binding protein, TEV cleavage sequence residues P6-P1, the relevant CTF sequence, and a FLAG tag | 114.6kDa | |
| HA-MBP-ADGRL2 CTFΔTA-FLAG | 832-1459 | pFuse-hIgG1-Fc2 | IL2 signal peptide followed by an HA tag, maltose binding protein, TEV cleavage sequence residues P6-P1, the relevant CTFΔTA sequence, and a FLAG tag | 113.9kDa | |
| HA-MBP-ADGRL3 CTF-FLAG | 842-1447 | pFuse-hIgG1-Fc2 | IL2 signal peptide followed by an HA tag, maltose binding protein, TEV cleavage sequence residues P6-P1, the relevant CTF sequence, and a FLAG tag | 111.4kDa | |
| HA-MBP-ADGRL3 CTFΔTA-FLAG | 849-1447 | pFuse-hIgG1-Fc2 | IL2 signal peptide followed by an HA tag, maltose binding protein, TEV cleavage sequence residues P6-P1, the relevant CTFΔTA sequence, and a FLAG tag | 110.7kDa | |
| HA-MBP-ADGRL4 CTF-FLAG | 407-690 | pFuse-hIgG1-Fc2 | IL2 signal peptide followed by an HA tag, maltose binding protein, TEV cleavage sequence residues P6-P1, the relevant CTF sequence, and a FLAG tag | 75.5kDa | |
| HA-MBP-ADGRL4 CTFΔTA-FLAG | 414-690 | pFuse-hIgG1-Fc2 | IL2 signal peptide followed by an HA tag, maltose binding protein, TEV cleavage sequence residues P6-P1, the relevant CTFΔTA sequence, and a FLAG tag | 74.8kDa | |
| HA-MBP-ADGRV1 CTF-FLAG | 5891-6306 | pFuse-hIgG1-Fc2 | IL2 signal peptide followed by an HA tag, maltose binding protein, TEV cleavage sequence residues P6-P1, the relevant CTF sequence, and a FLAG tag | 89.5kDa | |
| HA-MBP-ADGRV1 CTFΔTA-FLAG | 5898-6306 | pFuse-hIgG1-Fc2 | IL2 signal peptide followed by an HA tag, maltose binding protein, TEV cleavage sequence residues P6-P1, the relevant CTFΔTA sequence, and a FLAG tag | 88.8kDa | |
| Tricistronic Gs short TRUPATH | | pcDNA3.1 | Gαs short-Renilla, Gβ1, and GFP-γ2 are encoded on a single plasmid. Sequences encoding the Gαs short-Renilla and Gβ1 genes have a triple T2A sequence between them; the Gβ1 gene and GFP-γ2 are separated by an intervening IRES sequence. | | Justin English, Univ. Utah |
| Tricistronic Gq TRUPATH | | pcDNA3.1 | Gαq-Renilla, Gβ1, and GFP-γ2 are encoded on a single plasmid. Sequences encoding the Gαq-Renilla and Gβ1 genes have a triple T2A sequence between them; the Gβ1 gene and GFP-γ2 are separated by an intervening IRES sequence. | | Justin English, Univ. Utah |
| Tricistronic Gi1 TRUPATH | | pcDNA3.1 | Gαi1-Renilla, Gβ1, and GFP-γ2 are encoded on a single plasmid. Sequences encoding the Gαi1-Renilla and Gβ1 genes have a triple T2A sequence between them; the Gβ1 gene and GFP-γ2 are separated by an intervening IRES sequence. | | Justin English, Univ. Utah |
| Tricistronic G13 TRUPATH | | pcDNA3.1 | Gα13-Renilla, Gβ1, and GFP-γ2 are encoded on a single plasmid. Sequences encoding the Gα13-Renilla and Gβ1 genes have a triple T2A sequence between them; the Gβ1 gene and GFP-γ2 are separated by an intervening IRES sequence. | | Justin English, Univ. Utah |

| | | | | |
|---|---|---|---|---|
| HA-ADGRE5 FL | 20-835 | pFuse-hIgG1-Fc2 | IL2 signal peptide followed by an HA tag, full-length ADGRE5 isoform 1 | 90.9 kDa (unprocessed); 56.7 kDa (processed NTF) |
| HA-MBP-ADGRE5 CTF | 531-835 | pFuse-hIgG1-Fc2 | IL2 signal peptide followed by an HA tag, maltose binding protein, TEV cleavage sequence residues P6-P1, and the ADGRE5 CTF | 76.6 kDa |
| HA-MBP-ADGRE5 SL | 546-835 | pFuse-hIgG1-Fc2 | IL2 signal peptide followed by an HA tag, maltose binding protein, TEV cleavage sequence residues P6-P1, and ADGRE5 SL | 74.8 kDa |
| HA-BRIL-ADGRE5 CTF | 531-835 | pFuse-hIgG1-Fc2 | IL2 signal peptide followed by an HA tag, bone restricted ifitm-like protein (BRIL), TEV cleavage sequence residues P6-P1, and the ADGRE5 CTF | 48.0 kDa |
| HA-PGS-ADGRE5 CTF | 531-835 | pFuse-hIgG1-Fc2 | IL2 signal peptide followed by an HA tag, a glycogen synthase domain from Pyrococcus abyssi (PGS), TEV cleavage sequence residues P6-P1, and the ADGRE5 CTF | 58.1 kDa |
| HA-T4L-ADGRE5 CTF | 531-835 | pFuse-hIgG1-Fc2 | IL2 signal peptide followed by an HA tag, T4-lysozyme, TEV cleavage sequence residues P6-P1, and the ADGRE5 CTF | 54.4 kDa |
| HA-3xFLAG-ADGRE5 CTF | 531-835 | pFuse-hIgG1-Fc2 | IL2 signal peptide followed by an HA tag, a 3xFLAG tag, TEV cleavage sequence residues P6-P1, and the ADGRE5 CTF | 39.0 kDa |
| HA-MBP-S531 | 531-835 | pFuse-hIgG1-Fc2 | IL2 signal peptide followed by an HA tag, maltose binding protein, the CD97 CTF, and a FLAG tag | 75.8 kDa |
| HA-MBP-S532 | 532-835 | pFuse-hIgG1-Fc2 | IL2 signal peptide followed by an HA tag, maltose binding protein, an N-terminally-truncated CD97 CTF, and a FLAG tag | 75.7 kDa |
| HA-MBP-F533 | 533-835 | pFuse-hIgG1-Fc2 | IL2 signal peptide followed by an HA tag, maltose binding protein, an N-terminally-truncated CD97 CTF, and a FLAG tag | 75.6 kDa |
| HA-MBP-TEV-F533 | 533-835 | pFuse-hIgG1-Fc2 | IL2 signal peptide followed by an HA tag, maltose binding protein, TEV cleavage sequence residues P6-P1, an N-terminally truncated CD97 CTF, and a FLAG tag | 76.4 kDa |
| HA-MBP-ADGRE5 CTF S531A | 531-835 | pFuse-hIgG1-Fc2 | IL2 signal peptide followed by an HA tag, maltose binding protein, TEV cleavage sequence residues P6-P1, and the ADGRE5 CTF | 76.6 kDa |
| HA-MBP-ADGRE5 CTF S532A | 531-835 | pFuse-hIgG1-Fc2 | IL2 signal peptide followed by an HA tag, maltose binding protein, TEV cleavage sequence residues P6-P1, and the ADGRE5 CTF | 76.6 kDa |
| HA-MBP-ADGRE5 CTF F533A | 531-835 | pFuse-hIgG1-Fc2 | IL2 signal peptide followed by an HA tag, maltose binding protein, TEV cleavage sequence residues P6-P1, and the ADGRE5 CTF | 76.6 kDa |
| HA-MBP-ADGRE5 CTF A534G | 531-835 | pFuse-hIgG1-Fc2 | IL2 signal peptide followed by an HA tag, maltose binding protein, TEV cleavage sequence residues P6-P1, and the ADGRE5 CTF | 76.6 kDa |
| HA-MBP-ADGRE5 CTF I535A | 531-835 | pFuse-hIgG1-Fc2 | IL2 signal peptide followed by an HA tag, maltose binding protein, TEV cleavage sequence residues P6-P1, and the ADGRE5 CTF | 76.6 kDa |
| HA-MBP-ADGRE5 CTF L536A | 531-835 | pFuse-hIgG1-Fc2 | IL2 signal peptide followed by an HA tag, maltose binding protein, TEV cleavage sequence residues P6-P1, and the ADGRE5 CTF | 76.6 kDa |
| HA-MBP-ADGRE5 CTF M537A | 531-835 | pFuse-hIgG1-Fc2 | IL2 signal peptide followed by an HA tag, maltose binding protein, TEV cleavage sequence residues P6-P1, and the ADGRE5 CTF | 76.6 kDa |
| HA-MBP-ADGRE5 CTF A538G | 531-835 | pFuse-hIgG1-Fc2 | IL2 signal peptide followed by an HA tag, maltose binding protein, TEV cleavage sequence residues P6-P1, and the ADGRE5 CTF | 76.6 kDa |
| HA-MBP-ADGRE5 CTF H539A | 531-835 | pFuse-hIgG1-Fc2 | IL2 signal peptide followed by an HA tag, maltose binding protein, TEV cleavage sequence residues P6-P1, and the ADGRE5 CTF | 76.6 kDa |
| HA-MBP-ADGRE5 CTF S531K | 531-835 | pFuse-hIgG1-Fc2 | IL2 signal peptide followed by an HA tag, maltose binding protein, TEV cleavage sequence residues P6-P1, and the ADGRE5 CTF | 76.6 kDa |
| HA-MBP-ADGRE5 CTF S532K | 531-835 | pFuse-hIgG1-Fc2 | IL2 signal peptide followed by an HA tag, maltose binding protein, TEV cleavage sequence residues P6-P1, and the ADGRE5 CTF | 76.6 kDa |
| HA-MBP-ADGRE5 CTF F533K | 531-835 | pFuse-hIgG1-Fc2 | IL2 signal peptide followed by an HA tag, maltose binding protein, TEV cleavage sequence residues P6-P1, and the ADGRE5 CTF | 76.6 kDa |
| HA-MBP-ADGRE5 CTF A534K | 531-835 | pFuse-hIgG1-Fc2 | IL2 signal peptide followed by an HA tag, maltose binding protein, TEV cleavage sequence residues P6-P1, and the ADGRE5 CTF | 76.6 kDa |
| HA-MBP-ADGRE5 CTF I535K | 531-835 | pFuse-hIgG1-Fc2 | IL2 signal peptide followed by an HA tag, maltose binding protein, TEV cleavage sequence residues P6-P1, and the ADGRE5 CTF | 76.6 kDa |
| HA-MBP-ADGRE5 CTF L536K | 531-835 | pFuse-hIgG1-Fc2 | IL2 signal peptide followed by an HA tag, maltose binding protein, TEV cleavage sequence residues P6-P1, and the ADGRE5 CTF | 76.6 kDa |
| HA-MBP-ADGRE5 CTF M537K | 531-835 | pFuse-hIgG1-Fc2 | IL2 signal peptide followed by an HA tag, maltose binding protein, TEV cleavage sequence residues P6-P1, and the ADGRE5 CTF | 76.6 kDa |
| HA-MBP-ADGRE5 CTF A538K | 531-835 | pFuse-hIgG1-Fc2 | IL2 signal peptide followed by an HA tag, maltose binding protein, TEV cleavage sequence residues P6-P1, and the ADGRE5 CTF | 76.6 kDa |

| | | | | | |
|---|---|---|---|---|---|
| HA-MBP-ADGRE5 CTF H539K | 531-835 | pFuse-hIgG1-Fc2 | IL2 signal peptide followed by an HA tag, maltose binding protein, TEV cleavage sequence residues P6-P1, and the ADGRE5 CTF | 76.6 kDa | |
| HA-MBP-ADGRE5 CTF YDV->AAA | 531-835 | pFuse-hIgG1-Fc2 | IL2 signal peptide followed by an HA tag, maltose binding protein, TEV cleavage sequence residues P6-P1, and the ADGRE5 CTF | 76.6 kDa | |
| HA-MBP-ADGRE5 CTF EDW->AAA | 531-835 | pFuse-hIgG1-Fc2 | IL2 signal peptide followed by an HA tag, maltose binding protein, TEV cleavage sequence residues P6-P1, and the ADGRE5 CTF | 76.6 kDa | |
| secTEV Protease | | pcDNA3.1 | TEV protease variant engineered for activity in the secretory pathway | | Ref. 49 |
